# Learning with reward prediction errors in a model of the *Drosophila* mushroom body

**DOI:** 10.1101/776401

**Authors:** James E. M. Bennett, Andrew Philippides, Thomas Nowotny

**Affiliations:** Department of Informatics, University of Sussex, UK

**Author notes:** Corresponding author: James E. M. Bennett, School of Engineering and Informatics, University of Sussex, Chichester 1, Room 002, Falmer, Brighton, BN1 9QJ.

## Abstract

Effective decision making in a changing environment demands that accurate predictions are learned about decision outcomes. In *Drosophila*, such learning is orchestrated in part by the mushroom body (MB), where dopamine neurons (DANs) signal reinforcing stimuli to modulate plasticity presynaptic to MB output neurons (MBONs). Here, we extend previous MB models, in which DANs signal absolute rewards, proposing instead that DANs signal reward prediction errors (RPEs) by utilising feedback reward predictions from MBONs. We formulate plasticity rules that minimise RPEs, and use simulations to verify that MBONs learn accurate reward predictions. We postulate as yet unobserved connectivity, which not only overcomes limitations in the experimentally constrained model, but also explains additional experimental observations that connect MB physiology to learning. The original, experimentally constrained model and the augmented model capture a broad range of established fly behaviours, and together make five predictions that can be tested using established experimental methods.

## Introduction

Expedient decision making benefits from an organism’s ability to accurately predict the rewarding and punishing outcomes of each decision, so that it can meaningfully compare the available options and act to bring about the greatest reward. In many scenarios, an organism must learn to associate the valence of each outcome with the sensory cues predicting it. A broadly successful theory of reward learning is the delta rule (1), whereby reward predictions are updated in proportion to reward prediction errors (RPEs): the difference between predicted and received rewards. RPEs are more effective as a learning signal than absolute rewards because RPEs diminish as the prediction becomes more accurate, adding stability to the learning process. In mammals, RPEs are signalled by dopamine neurons (DANs) in the ventral tegmental area and substantia nigra, enabling the brain to implement approximations to the delta rule (2; 3). In *Drosophila melanogaster*, DANs that project to the mushroom body (MB) (Fig. 1a) also provide reward modulated signals that are required for associative learning (4). However, to date, MB DAN activity is typically interpreted as signalling absolute reinforcement, and direct evidence for RPE signals is lacking. Here, we build a novel computational model of the MB, taking advantage of precise anatomical and functional data from recent experiments, and provide a circuit-level description for delta rule learning in the MB, whereby MB DANs compute RPEs.

**Figure 1:**
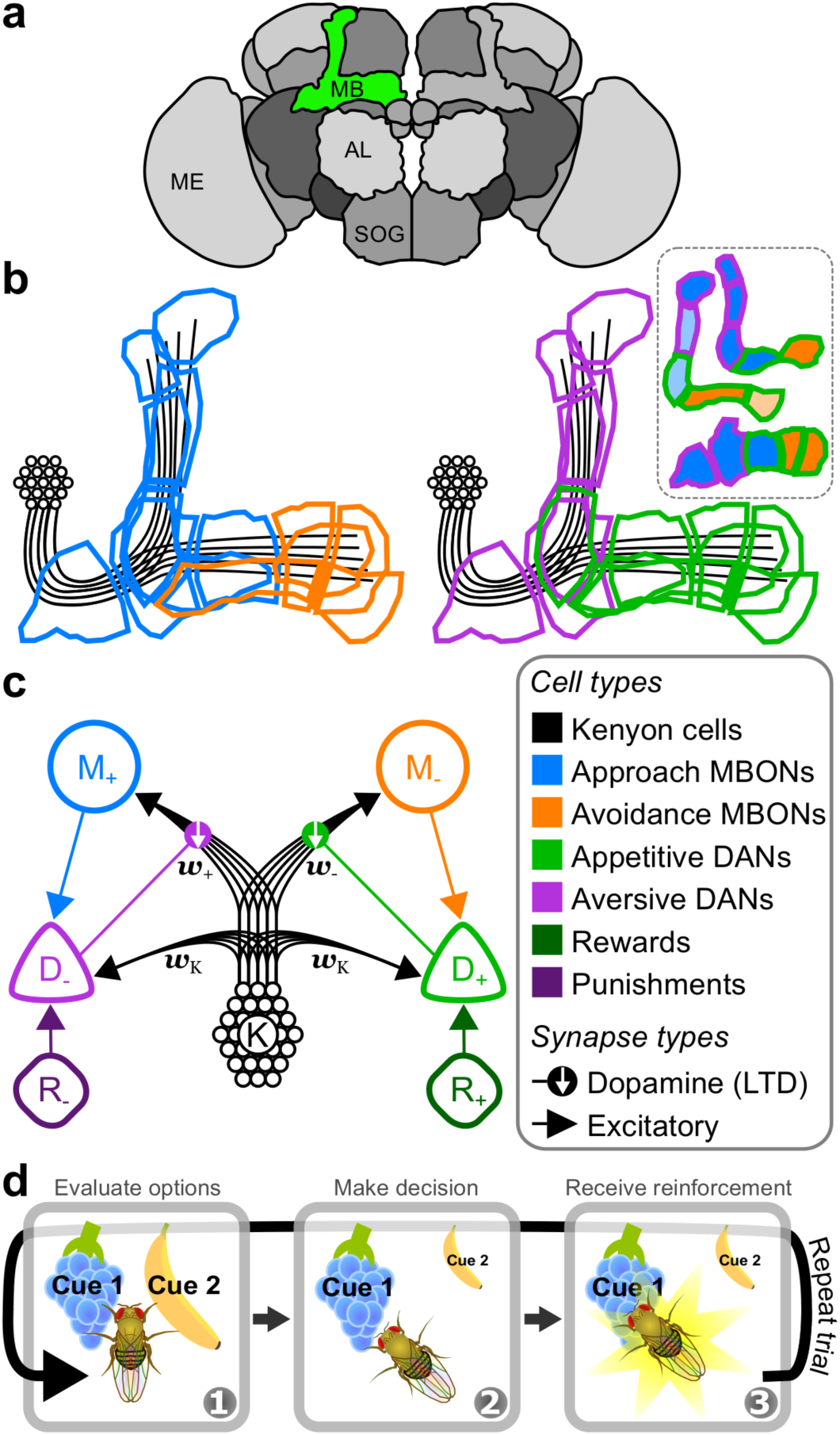
Valence-specific model of the mushroom body. **a** 3D render of several neuropils that comprise the brain of *Drosophila melanogaster*. The red region highlights the MB in the right hemisphere. Labels: MB – mushroom body; AL – antennal lobe; SOG – suboesophogeal ganglion; ME – medulla. **b** Outlines of the multiple compartments that tile the lobes of the MB, colour-coded by a broad classification of cell function. Blue: approach MBONs; orange: avoidance MBONs; purple: aversive DANs; green: appetitive DANs; black: KCs. *Inset*: schematic of the three MB lobes and their compartmentalisation. Top: *α*′/*β*′ lobes; middle: *α*/*β* lobes; bottom: *γ* lobe. MBON functions (approach or avoidance) are as determined in (9). Pale colours in the *inset* correspond to MBONs that exhibit a non-significant bias on behaviour in (9). **c** Schematic of the VS model. Units are colour-coded according to the cell types in **b**. KCs connect to MBONs through plasticity synapses, and connect to DANs through fixed synapses. Labels: M_+_ – approach MBON; M_−_ – avoidance MBON; D_−_ – aversive DAN; *D*_+_ – appetitive DAN; K – Kenyon cells; R_−_ – negative reward; R_+_ – positive reward. Lines with arrows: excitatory synapse. Lines with filled circles: synapse releasing dopamine. Downward white arrows: dopamine enhances synaptic depression. **d** Schematic of a single trial for the experimental paradigm in which the model is examined. Panel 1: the model is exposed to some number of cues that are evaluated to yield a cue-specific reward prediction. Panel 2: using the relative reward predictions, the model makes a decision over which cue to choose. Panel 3: the model receives a reinforcement signal that is associated with the chosen cue, and its reward predictions for that cue are updated.

Consistent with its role in associative learning, the MB is organised into lateral and medial lobes of neuropil that, in addition to DANs, are innervated by the axons of sensory encoding Kenyon cells (KCs), and the dendrites of MB output neurons (MBONs), which modulate behaviour (Fig. 1b). Current models of MB function posit that the MB lobes encode different valences of reward information and actions (5–11). Most DANs in the PAM (protocerebral anterior medial) cluster (called D_+_ in the model presented here) are activated by rewards, or positive reinforcement (R_+_), and their activation results in depression at synapses between coactive KCs (K) and MBONs that are thought to induce avoidance behaviours (M_−_). DANs in the PPL1 (protocerebral posterior lateral 1) cluster (D_−_) are activated by punishments, i.e. negative reinforcement (R_−_), and their activation results in depression at synapses between coactive KCs and MBONs that induce approach behaviours (M_+_). A fly can therefore learn to approach rewarding cues or avoid punishing cues as a result of synaptic depression at KC inputs to avoidance or approach MBONs, respectively.

To date, there are only indirect lines of evidence for RPE signals in MB DANs. DAN activity is modulated by feedforward reward signals, but some DANs also receive excitatory feedback from MBONs (12–15), and it is likely this extends to all MBONs whose axons are proximal to DAN dendrites (16). MBON activity is, in turn, modulated by DANs via plasticity at KC→MBON synapses (10; 17; 18). We interpret the difference between approach and avoidance MBON firing rates as a reward prediction that motivates behaviour, consistent with the observation that behavioural valence scales with the difference between approach and avoidance MBON firing rates (9). As such, DANs that integrate feedforward reward signals and feedback reward predictions from MBONs are primed to signal RPEs for learning. To the best of our knowledge, these latter two features have yet to be incorporated in computational models of the MB (19–21).

Here, we formulate a reduced computational model of the MB circuitry described above to demonstrate how DANs may compute RPEs, then, hypothesising that learning minimises RPEs, derive a plasticity rule for KC→MBON synapses, and verify in simulations that it yields accurate reward predictions. We identify limitations to the model that impose an upper bound to reward prediction magnitudes, and demonstrate how as yet unobserved connections between DANs, KCs and MBONs could circumvent this limitation. Introducing these additional connections yields testable predictions for future experiments as well as explaining a broader range of existing experimental observations that connect DAN and MBON stimulus responses to learning. Lastly, we show that both incarnations of the model – with and without additional connections – capture a wide range of experimental observations in *Drosophila*, whereby conditioned approach and avoidance behaviours are compromised by genetically targeted manipulation of MBON and DAN activity. Different behavioural outcomes in the two models for particular genetic manipulations provide further strong experimental predictions.

## Results

### A computational model of the mushroom body

The MB lobes comprise multiple compartments, each innervated by a different set of MBONs and DANs (Fig. 1b), and each encoding memories for different forms of reward (22), with different longevities (23), and for different stages of memory formation (24). Nevertheless, compartments appear to contribute to learning by similar mechanisms (10; 18; 25), and it is reasonable to assume that the process of learning reward predictions is similar for different forms of reward. We therefore reduce the multicompartmental MB into two compartments, and assign a single, rate-based unit to each class of MBON and DAN (colour-coded in Fig. 1B-C). KCs, however, are modelled as a population, in which each sensory cue selectively activates a unique subset of 10 units. Given that activity in approach and avoidance MBONs – denoted M_+_ and M_−_ in our model – respectively bias flies to approach or avoid a cue, *i*, we interpret the difference in their firing rates, 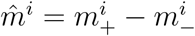, as the fly’s reward prediction for that cue.

For the purpose of this work, we assume that the MB has only a single objective: to form reward predictions that are as accurate as possible, i.e. that minimise the RPE. We do this within a multiple-alternative forced choice (MAFC) paradigm (Fig. 1D; also known as a multi-armed bandit) in which a fly is exposed to one or more sensory cues in a given trial, and is forced to choose one. The fly then receives a reinforcement signal, 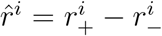, which has both positive and negative components (coming from sources R_+_ and R_−_), and which is specific to the chosen cue. Over several trials, the fly must learn to predict the rewards for each cue, and use these predictions to reliably choose the most rewarding cue. We formalise this objective with a cost function

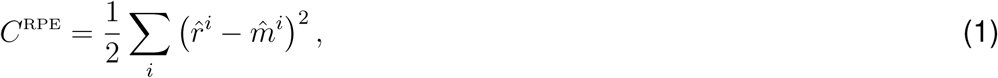

where the sum is over all cues, *i*. By performing gradient descent on *C*^RPE^, we derived a learning rule, *𝒫*^RPE^ (full derivation in Methods: Synaptic plasticity):

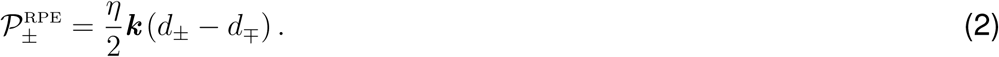

which minimises the cost when used to update synaptic weights according to 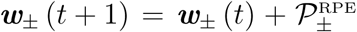. We use subscripts ‘±’ to denote the valence of the neuron: if + (−) is considered in ±, then ∓ refers to − (+), and *vice versa*. As such, *d*_±_ refers to the firing rate of either D_+_ or D_−_. The learning rate, *η*, must be small (see Methods: Synaptic plasticity) to allow the plasticity rule to average over multiple stimuli as well as stochasticity in the reward schedule (see Methods: Reward schedule). Note that a single DAN, D_±_, only has access to half of the reward and reward prediction information, and by itself does not compute the full RPE. However, the difference between D_+_ and D_−_ firing rates does yield the full RPE (see Methods: DAN firing rates):

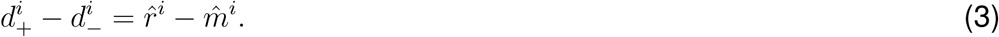

Three features of Eq. 2 are worth highlighting here. First, elevations in *d*_±_ increase the net amount of synaptic depression at active synapses that impinge on M_∓_, which encodes the opposite valence to D_±_, in agreement with experimental data (10; 18; 25). Second, the postsynaptic MBON firing rate is not a factor in the plasticity rule, unlike in reward-modulated Hebbian rules (26), yet nevertheless in accordance with experiments (18). Third, Eq. 2 requires that synapses receive dopamine signals from both D_+_ and D_−_, conflicting with current experimental findings in which appetitive DANs only modulate plasticity at avoidance MBONS, and similarly for aversive DANs and approach MBONs (10; 17; 18; 22; 27; 28). To accommodate the experimental data, we consider an alternative cost function, 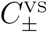, that satisfies the valence specificity of DANs:

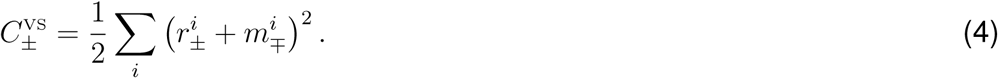

Gradient descent on 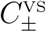 yields the corresponding valence-specific plasticity rule:

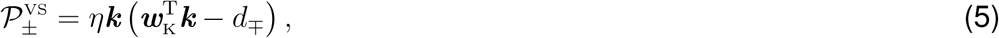

where 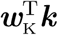 models the direct excitatory current from KCs to DANs (Methods, Eq. 11). As required, Eq. 5 maintains the relationship between increased DAN activity and enhanced synaptic depression. We refer to models that utilise this plasticity rule as valence-specific (VS) models. In our simulations, we update only those synaptic weights that correspond to the chosen cue, as only the reward for this cue would be experienced by a fly (see Methods, Eq. 21).

### Restricted learning in the valence-specific model

Eq. 5 exposes a problem for learning according to our assumed objective in the VS model. The problem arises because D_±_ receives only excitatory inputs. Thus, whenever a cue is present, KC inputs (29) prescribe D_±_ with a minimum, cue-specific firing rate, 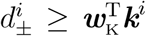. As such, synapses will be depressed 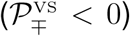 whenever *r*_±_ > 0 or *m*_∓_ > 0. Once 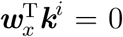, the VS model can no longer learn the valence of cue *i* as synaptic weights cannot become negative. Eventually, reward predictions for all cues become equal with 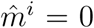, such that choices become random (Supplementary Fig. 1a-b). In this case, D_+_ and D_−_ firing rates become equal to the positive and negative rewards, respectively, such that RPEs equal the net rewards (Supplementary Fig. 1c-d).

A heuristic solution is to add a constant source of potentiation, which acts to restore synaptic weights to a constant, non-zero value. We therefore replace 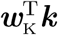 in Eq. 5 with a constant, free parameter, *λ*:

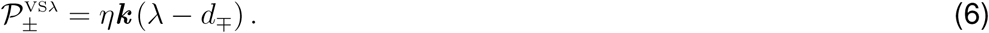

By setting 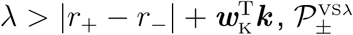 can take both positive and negative values, preventing synaptic weights from being held at zero. This defines a new baseline firing rate for D_±_ that is greater than 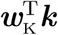. Hereafter, we refer to the VS model with plasticity governed by 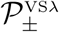 as the VS*λ* model.

The VS*λ* model provides only a partial solution, as it is restricted by an upper bound to the magnitude of reward predictions that can be learned: 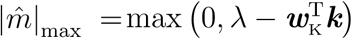. In addition to increasing *λ*, 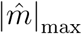 may be increased by reducing KC→DAN synaptic transmission. In Fig. 2a, we set ***w***_K_ = *γ***1**, with **1** a vector of ones, and show reward predictions for several values of *γ*, with *λ* = 11.5 (corresponding DAN and MBON firing rates are in Supplementary Fig. 2). The upper bound is reached when ***w***_+_ or ***w***_−_, and thus the corresponding MBON firing rates, go to zero (an example when *γ* = 1 is shown in Fig. 2b). These results appear to contradict recent experimental work in which learning was impaired, rather than enhanced, by blocking KC→DAN synaptic transmission (29).

**Figure 2:**
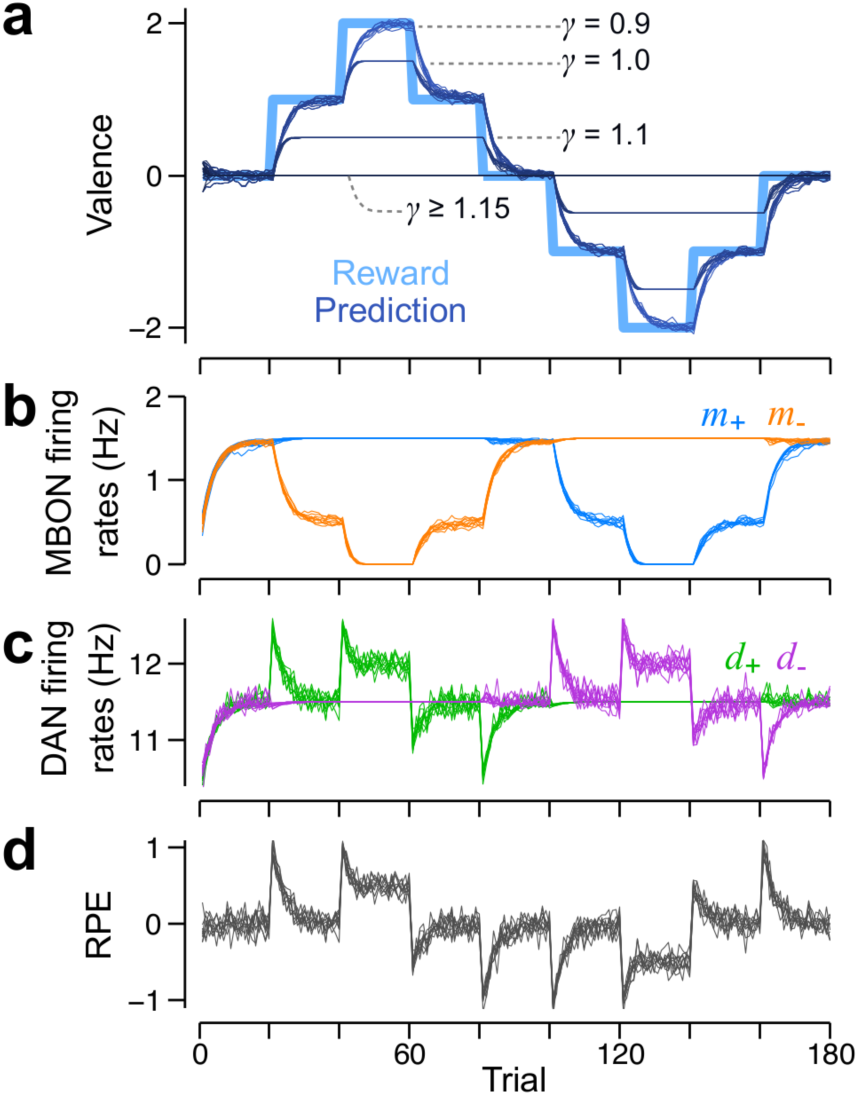
Reward predictions in the valence-specific model track rewards but only within specified bounds. **a** Reward schedule (excluding the Gaussian white noise; thick, light blue) and reward predictions (thin, various dark blues) from 10 separate runs of the model. Each shade of dark blue corresponds to simulations using a specific KC→DAN synaptic weight, which is determined by *γ*: dark blue through to very dark blue corresponds to *γ* = 0.9, 1.0, 1.1, and *γ* > 1.15. Dashed lines correspond to 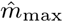, the theoretical maximum to the reward prediction magnitude, for each value of *γ*. **B** Firing rates of the M_+_ (blue) and the M_−_ MBONs (orange) for 10 runs of the model, with the same reward schedule as in **a. c** Firing rates for the D_+_ (green) and the D_−_ (purple) DANs in response to the reward schedule in **a. d** RPEs given by the difference in firing rates of D_+_ and D_−_.

In the VS*λ* model, DAN firing rates begin to exhibit RPE signals. A sudden increase in positive reinforcements, for example at trial 20 in Fig. 2c, results in a sudden increase in *d*_+_, which then decays as the excitatory feedback from M_−_ diminishes as a result of synaptic depression in ***w***_−_ (Fig. 2c-d). Similarly, sudden decrements in positive reinforcements, for example at trial 80, are signalled by reductions in *d*_+_. However, when the reward magnitude exceeds the upper bound, as in trials 40-60 and 120-140 in Fig. 2a-d, D_±_ exhibits sustained elevations in firing rate from baseline by an amount 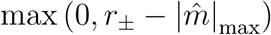 (Fig. 2c, Supplementary Fig. 2). This constitutes a major prediction from our model.

### A mushroom body circuit with unbounded learning

In the VS*λ* model, excitatory reward signals can only be partially offset by decrements to ***w***_+_ and ***w***_−_. Thus, to overcome the upper bound to reward predictions, a source of inhibition is required. A candidate solution is a circuit in which positive reinforcements, R_+_, inhibit D_−_, and similarly, R_−_ inhibits D_+_ (illustrated in Fig. 3a). Such inhibitory reinforcement signals have been observed in the *γ*2, *γ*3, *γ*4 and *γ*5 compartments of the MB (17; 30). Using the derived plasticity rule, 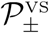 in Eq. 5, this circuit learns accurate reward predictions with no upper bound to the reward prediction magnitude (Supplementary Fig. 3a). Hereafter, we refer to the VS model with unbounded learning as the VSu model. Learning is now possible because, when the synaptic weights ***w***_±_ are weak, or when D_∓_ is inhibited, Eq. 5 specifies that 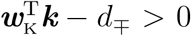, i.e. synaptic weights will potentiate until the feedback excitation equals the reinforcement-induced feedforward inhibition. Similarly, synapses are depressed in the absence of reinforcement because the excitatory feedback from M_±_ to D_∓_ ensures that 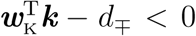 (Supplementary Fig. 3b). Consequently, step changes in reinforcement yield RPE signals in D_∓_ that always decay to a baseline set by 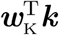 (Supplementary Fig. 3c-d). Despite the prevalence in reports of long term synaptic depression in the MB, there exist several lines of evidence for potentiation as well (10; 11; 14; 31). However, when reinforcement signals are inhibitory, D_+_, for example, is activated only after decrements to *r*_−_, but not following increments to *r*_+_ (similarly for D_−_), counter to the experimental classification of DANs as appetitive (or aversive) (6–8; 32).

**Figure 3:**
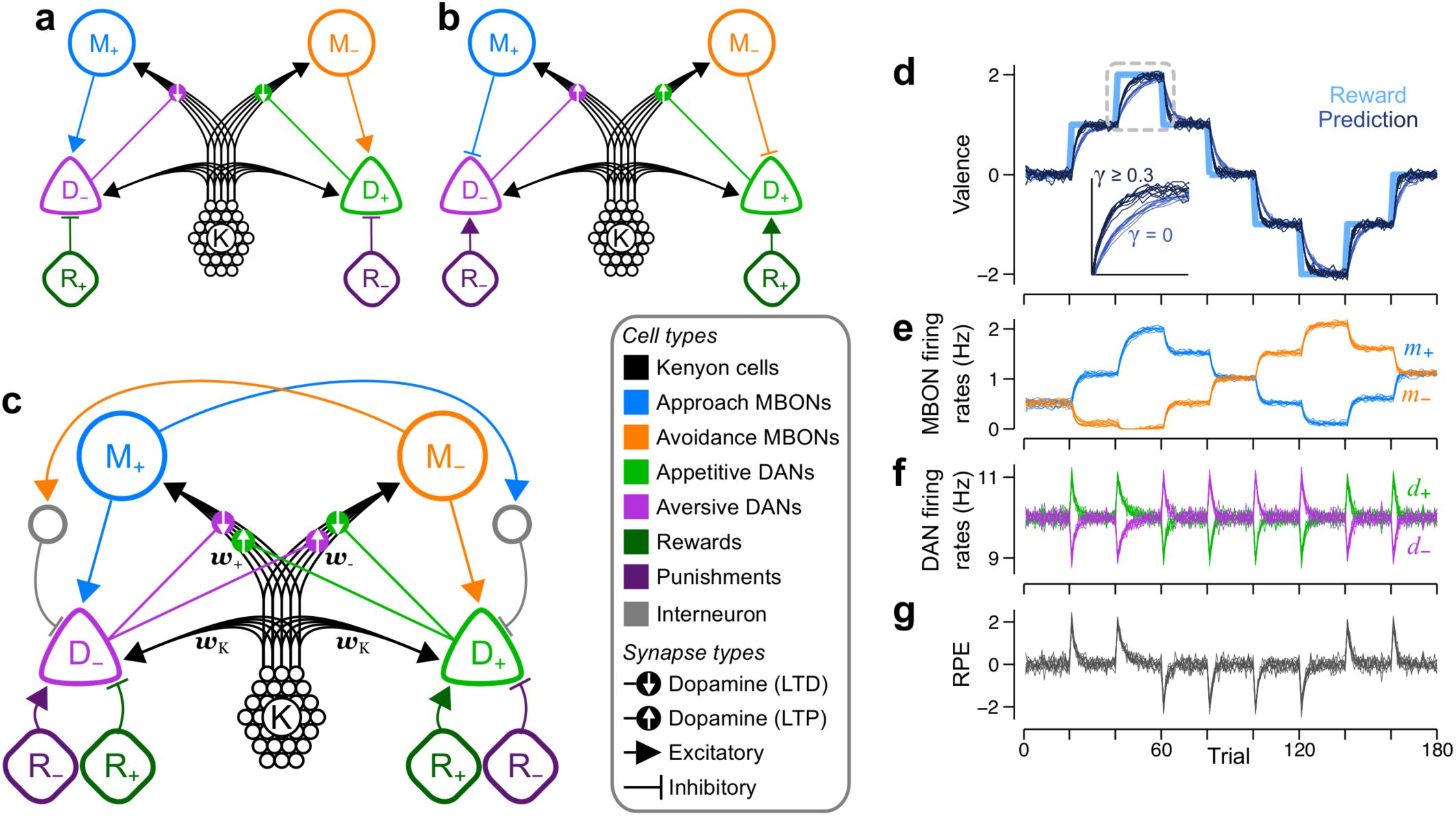
Dual versions of unbounded valence-specific (VSu) models can be combined to create the mixed-valence (MV) model, which also learns unbounded reward predictions. **a** One of the dual, VSu models that requires D_−_ and *D*_+_to be inhibited by positive and negative rewards, respectively. Lines with flat ends correspond to inhibitory synapses. **b** The second of the dual VSu models, in which MBONs provide inhibitory feedback to DANs of the same valence. Right, upward arrows in the dopamine synapses denote that dopamine induces synaptic potentiation. **c** The MV model, which combines the dual VSu models. Grey units are inhibitory interneurons. **d**-**e** Each panel exhibits the behaviour from 10 independent runs of the model. **d** Reward predictions are unbounded and accurately track the rewards, but the learning speed depends on KC→DAN synaptic weights. Thick, light blue: reward schedule; thin, dark blue: *γ* = 0; thin, very dark blue: *γ ≥* 0.3. *Inset*: magnified view of the region highlighted by the dashed square, showing how learning is slower when *γ* = 0. **e** M_+_ (blue) and M_−_ firing rates when *γ* = 1. **f** D_+_ and D_−_ firing rates when *γ* = 1. **g** RPEs as given by the difference between D_+_ and D_−_ firing rates when *γ* = 1.

We therefore asked whether other variations of the VSu model could learn without an upper bound, and identified three criteria (tabulated in Supplementary Table 1) that must be satisfied to achieve this: *i*) learning must be effective, such that positive reinforcement either potentiates excitation of approach behaviours (inhibition of avoidance), or depresses inhibition of approach behaviours (excitation of avoidance), and similarly for negative reinforcement; *ii*) learning must be stable, such that excitatory reinforcement signals are offset via learning, either by synaptic depression of feedback excitation, or by potentiation of feedback inhibition, and similarly for inhibitory reinforcement signals; *iii*) to be unbounded, learning must involve synaptic potentiation, whether reinforcement signals excite DANs that induce potentiation, or inhibit DANs that induce depression. By following these criteria, we identified a dual version of the VSu circuit in Fig. 3a, which is illustrated in Fig. 3b. In this circuit, R_+_ excites D_+_ and R_−_ excites D_−_. However, DANs induce synaptic potentiation when activated above baseline, while M_+_ and M_−_ are inhibitory, so are interpreted as inducing avoidance and approach behaviours, respectively. Despite their different configurations, reward predictions are identical in each of the dual MB circuits (data not shown).

Neither dual model, by itself, captures all of the experimentally established anatomical and physiological properties of the MB. However, by combing them into one (Fig. 3c), the new model becomes consistent with the circuit properties observed in experiments, but necessitates additional features, which constitute major predictions. First, synapses impinging on either approach or avoidance MBONs must undergo plasticity that is modulated by DANs of both valences. Second, DANs receive both positive and negative reward signals, which are either excitatory or inhibitory, depending on the valences of the reinforcement and the DAN. Third, MBONs provide feedback to DANs of the same valence via inhibitory interneurons, which we propose innervate areas targeted by MBON axons and DAN dendrites (16). Each DAN in this hybrid model now has access to the full reward, 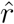, and the full reward prediction, 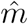, or 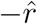 and 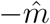, depending on the valence of the DAN. Thus, it is now reasonable to utilise the original plasticity rule,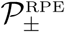 in Eq. 2, which was derived with the objective to minimise the full RPE. As such, DANs facilitate synaptic depression if the DAN and MBON have opposite valences, or facilitate potentiation if they have the same valence. We refer to this new model, under the control of 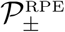, as the mixed valence (MV) model.

Fig. 3d demonstrates that the MV model accurately tracks changing rewards, just as with the dual versions of the VSu model. However, a number of differences from the VSu models can also be seen. First, changing reward predictions are manifest by changes in the firing rates of both M_+_ and M_−_ (Fig. 3e). Moreover, when ***w***_±_ reach zero, the changes in ***w***_∓_ compensate, resulting in larger changes in the firing rate of M_∓_, as seen between trials 40-60 in Fig. 3e. Second, DANs no longer encode prediction errors exclusively in either positive or negative reinforcers, but their activity is instead correlated with the sign of the full RPE: *d*_+_ and *d*_−_ increase with positive and negative RPEs, respectively, and decrease with negative and positive RPEs (Fig. 3f-g). Third, blocking KC→DAN synaptic transmission slows down learning, but does not abolish it entirely (Fig. 3d). With input from KCs blocked, the baseline firing rate of D_±_ is zero, and because any given RPE excites one DAN type and inhibits the other, only one of either D_+_ or D_−_ can signal the RPE, reducing the magnitude of *d*_±_ − *d*_∓_ in Eq. 2, and therefore the speed of learning (Supplementary Fig. 4). To avoid any slowing down to learning, 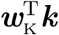 must be greater than or equal to the RPE. This may explain the 25% reduction in learning performance seen by Cervantes-Sandoval et al. (29).

### Decision making in a multiple-alternative forced choice task

We next tested the VS*λ* and MV models on a task with multiple cues from which to choose. Choices are made using the softmax function (Eq. 9), such that the model more reliably chooses one cue over another when cue-specific reward predictions are more dissimilar. Throughout the task, the cue-specific reinforcements slowly change (see example reward schedules in Fig. 4a-b), and the model must continually update reward predictions, according to its plasticity rule, in order to choose the most positively reinforcing cues as possible.

**Figure 4:**
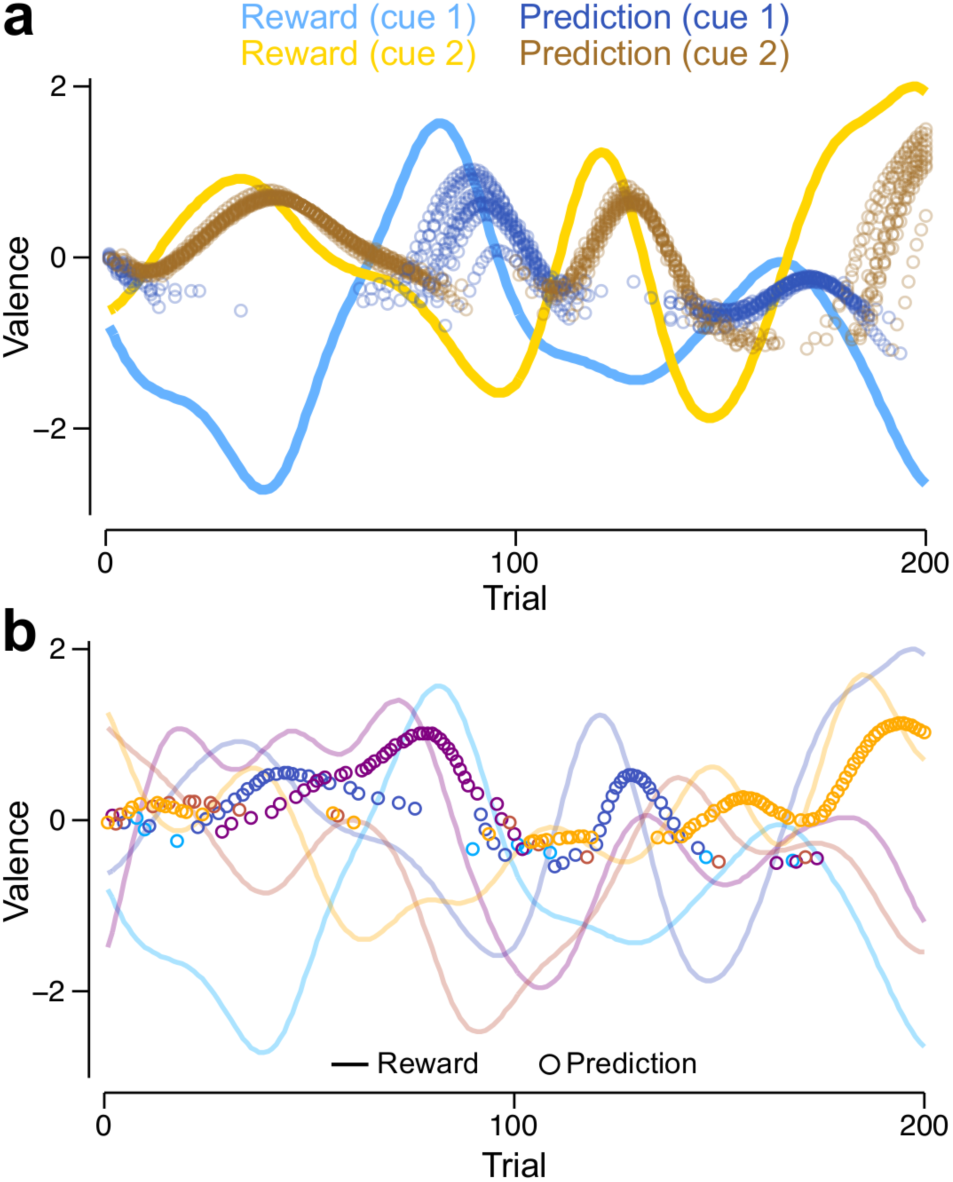
Learning reward predictions in tasks with multiple cues. **a** Reward schedules (lines) and reward predictions (circles, shown only for the cue chosen on each trial) for two cues (blue: cue 1; yellow: cue 2). Reward predictions are shown for 10 independent runs of a simulation using the same reward schedule. **b** Reward schedules (lines) and reward predictions (circles, shown only for the cue chosen on each trial) for a single run of the model in a task involving 5 cues. Each colour corresponds to rewards and predictions for a different cue.

In a task with two alternatives, switches in cue choice almost always occur after the actual switch in the reinforcement schedule because of the slow learning rate and the probabilistic nature of decision making (Fig. 4a). The model continues to choose the more rewarding cues when there are as many as 200 (Supplementary Fig. 5a; Fig. 4b shows an example simulation with 5 cues). When there are few cues, the mean reward per trial (RPT) obtained increases with the number of cues, as the number of cues providing a large reward increases. The RPT then decreases monotonically with the number of cues as the model is less able to maintain accurate predictions for all cues. Despite this latter degradation in performance, the VS*λ* and MV models are only marginally outperformed by a model with perfect plasticity, whereby reward predictions for the chosen cue are set to equal the last obtained reward. Furthermore, when noise is added to the reinforcement schedule, the performance of the perfect plasticity model drops below that of the other models, for which slow learning helps to average over noisy observations (Supplementary Fig. 5b). The model suffers no noticeable decrement in performance when KC responses to different cues overlap, e.g. when a random 5% of 2000 KCs are assigned to each cue (Supplementary Fig. 5a,c-e).

### Both models capture learned fly behaviours in a variety of conditioning experiments

To determine how well the VS*λ* and the MV models capture decision making in flies, we applied them to an experimental paradigm (illustrated in Fig. 5a) in which flies are conditioned to approach or avoid one of two odours. In each experiment, flies undergo a training stage, during which they are exposed to a conditioned stimulus (CS+) concomitantly with an unconditioned stimulus (US), for example sugar (appetitive training) or electric shock (aversive training). Flies are next exposed to a different stimulus (CS-) without any US. Following training, flies are tested for their behavioural valence with respect to the two odours. The CS+ and CS- are released at opposite ends of a tube. Flies are free to approach or avoid the stimuli by walking towards one end of the tube or the other. In our model, we do not simulate the spatial extent of the tube, nor specific fly actions, but model choice behaviour in a simple manner by applying the softmax function to the current reward predictions.

**Figure 5:**
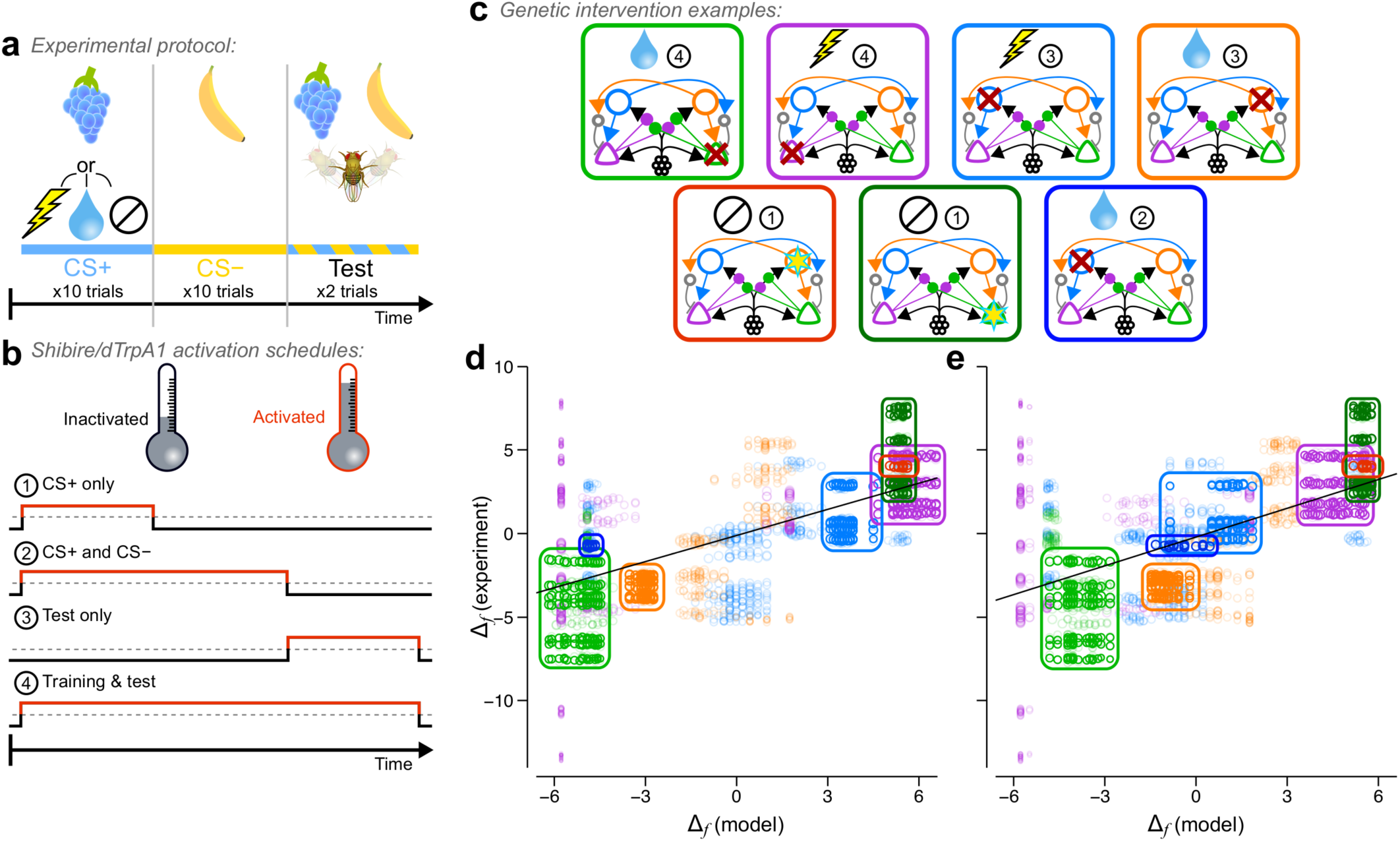
Both the modified valence-specific (VS*λ*) and mixed valence (MV) models produce choice behaviour that corresponds well with experiments under a broad range of experimental manipulations. **a**) Schematic of the experimental protocol used to simulate appetitive, aversive and neutral conditioning experiments. **b-c**) The protocol was extended to simulate genetic interventions used in experiments. **b**) Interventions were applied at different stages of a simulation, either 1) during CS+ exposure in training, 2) during CS+ and CS− exposure in training, 3) during testing, and 4) throughout both training and testing. **c**) Four examples of the interventions simulated, each corresponding to encircled data in **d** and **e**. Red crosses denote interventions that simulate activation of a *shibire* blockade; yellow stars denote interventions that simulate activation of an excitatory current through the dTrpA1 channel. The picture at the top of each panel denotes the reward type, and the encircled number the activation schedule as specified in **b. d**) Comparison of PIs measured from the VS*λ* model and from experiments. **e**) Comparison of PIs measured from the MV model and from experiments. Grey lines in **d** and **e** are weighted least square linear fits with correlation coefficients *R* = 0.65 and *R* = 0.63 respectively. The size of each data point scales with its weight in the linear fit.

In addition to these control experiments, we simulated a variety of interventions frequently used in experiments (Fig. 5a-c). These experiments are determined by four features: 1) US valence (Fig. 5a): appetitive, aversive, or neutral; 2) intervention type (Fig. 5c): inhibition of neuronal output, e.g. by expression of *shibire*, or activation, e.g. by expression of dTrpA1, both of which are controlled by temperature; 3) the intervention schedule (Fig. 5b): during the CS+ only, throughout CS+ and CS−, during test only, or throughout all stages; 4) the target neuron (Fig. 5c): either M_+_, M_−_, D_+_, or D_−_. Further details of these simulations are provided in Methods: Experimental data and model comparisons.

We compared the models to behavioural results from 439 experiments (including 235 controls), which tested 27 unique combinations of the above four parameters in 14 previous studies (7–13; 22; 23; 27; 30; 31; 33; 34) (data provided in Supplementary Table 2). The source and experimental details of each experimental intervention used here is listed in Supplementary Table 2. In Fig. 5d-e, we plot a test statistic, Δ_*f*_, that compares behavioural performance indices (PIs) between a specific intervention experiment and its corresponding control, where the PI is +1 if all flies approached the CS+, and −1 if all flies approached the CS−. For example, a positive Δ_*f*_ indicates that more flies approached the CS+ in the intervention than in the control experiment. Interventions in both models correspond well with those in the experiments: Δ_*f*_ from the VS*λ* model and experiments are correlated with *R* = 0.65, and Δ_*f*_ from the MV model and experiments are correlated with *R* = 0.63 (*p <* 0.0001 for both models).

Four cases of inhibitory interventions exemplify the correspondence of both the VS*λ* and MV model with experiments, and are highlighted in Fig. 5d-e (light green, purple, blue and orange rings). Also highlighted are two examples of excitatory interventions, in which artificial stimulation of either D_+_ or M_−_ during CS+ exposure, without any US, was used to induce an appetitive memory and approach behaviour. The two models do not always yield the same behavioural result. The example highlighted in dark blue, in which M_+_ was inhibited throughout appetitive training but not during the test, shows that this intervention had little effect in the MV model, in agreement with experiments (31), but resulted in a strong reduction in the appetitiveness of the CS+ in the VS*λ* model (Δ_*f*_ ≈ −4.5). In the Supplementary Text, we analyse the underlying synaptic weight dynamics that lead to this difference in model behaviours. The analyses show that not only does this intervention amplify the difference between CS+ and CS− reward predictions in the MV model, it also results in faster memory decay in the VS*λ* model. Hence, the preference for the CS+ is maintained in the MV model, but is diminished in the VS*λ* model.

## Discussion

### Overview

Successful decision making relies on the ability to accurately predict, and thus reliably compare, the outcomes of choices that are available to an agent. The delta rule, developed by Rescorla & Wagner (1), which updates beliefs in proportion to a prediction error, provides a method to learn accurate predictions. In this work, we have investigated the hypothesis that, in *Drosophila melanogaster*, the MB implements the delta rule. We posit that approach and avoidance MBONs together encode reward predictions, and that feedback from MBONs to DANs, if subtracted from feedforward reinforcement signals, endows DANs with the ability to compute RPEs, which are used to modulate synaptic plasticity. We formulated a plasticity rule that minimises RPEs, and verified the effectiveness of the rule in simulations of multiple-alternative forced choice tasks. We demonstrated how the established valence-specific circuitry of the MB restricted the learned reward predictions to within a given range, and postulated additional anatomical features that could overcome this restriction. We have thus presented two MB models that yield RPEs in DAN activity and that learn accurate reward predictions: *i*) the VS*λ* model, in which plasticity incorporates a constant source of synaptic potentiation; *ii*) the MV model, in which we propose mixed-valence connectivity between DANs, MBONs and KC→MBON synapses. Both the VS*λ* and the MV models receive equally good support from behavioural experiments in which different genetic interventions impaired learning, while the MV model provides a mechanistic account for a greater variety of physiological changes that occur in individual neurons after learning. It is plausible, and can be beneficial, for both the VS*λ* and MV models operate in parallel in the MB, as separately learning positive and negative aspects of decision outcomes, if they arise from independent sources, is important for context-dependent modulation of behaviour. Such learning has been proposed for the mammalian basal ganglia (35).

### Predictions

The models yield predictions that can be tested using established experimental protocols. Below, we specify which model supports each prediction.

#### Prediction 1 – both models

DAN responses to the unconditioned stimulus (US) should decay towards a baseline over successive CS+-US pairings, as a result of the learned changes in MBON firing rates. To the best of our knowledge, only one study has measured DAN responses throughout learning in *Drosophila* (36). Consistent with DAN responses in our model, Dylla et al. (36) reported such decaying responses in DANs in the *γ*- and *β*′-lobes during paired CS+ and US stimulation. However, they reported similar decaying responses when the CS+ and US were unpaired (separated by 90 seconds) that were not significantly different from the paired condition. The authors concluded that DANs do not exhibit RPEs, and that the decaying DAN responses were a result of non-associative plasticity. An alternative interpretation is that a 90 second gap between CS+ and US does not induce DAN responses that are significantly different from the paired condition, and that additional processes prevent the behavioural expression of learning. Ultimately, the evidence for either effect is insufficient. Moreover, Dylla et al. (36) observed increases in DAN responses to the CS+ during training. This is consistent with the temporal difference learning algorithm (37; 38), which is an extension of the Rescorla-Wagner rule.

#### Prediction 2 – VSλ model

a sufficiently large reward will yield a sustained DAN response. This feature may be tested by conditioning *Drosophilae* with a stepped electric shock schedule. DAN responses may initially exhibit RPEs, which should transition to sustained responses as the voltage increases. The absence of sustained responses could imply that *λ* adapts to the reward history.

#### Prediction 3 – MV model

the valence of a DAN is defined by its response to RPEs, rather than to rewards *per se*. Thus, DANs previously thought to be excited by positive (negative) reinforcement are in fact excited by positive (negative) RPEs. For example, a reduction in electric shock magnitude, after an initial period of training, would elicit an excitatory (inhibitory) response in appetitive (aversive) DANs. Felsenberg et al. (13; 14) provide indirect evidence for this. The authors trained flies on a CS+, then re-exposed the fly to the CS+ without the US. For an appetitive (aversive) US, CS+ re-exposure would have yielded a negative (positive) RPE. By blocking synaptic transmission from aversive (appetitive) DANs during CS+ re-exposure, the authors prevented the extinction of learned approach (avoidance).

#### Prediction 4 – MV model

learning is mediated by simultaneous plasticity at both approach and avoidance MBON inputs. Appetitive conditioning does indeed potentiate responses in MB-V3/*α*3 and MVP2/*γ*1-pedc approach MBONs (11; 31), and depress responses in M4*β*′/*β*′2mp and M6/*γ*5*β*′2a avoidance MBONs (10). Similarly, removal of an expected aversive stimulus, which constitutes a positive RPE, depresses M6/*γ*5*β*′2a avoidance MBONs (14). In addition, aversive conditioning depresses responses in MPV2/*γ*1-pedc MB-V2/*α*2*α*′2 approach MBONs (18; 25), and potentiates responses in M4*β*′/*β*′2mp and M6/*γ*5*β*′2a avoidance MBONs (10; 24). However, the potentiation of M4*β*′ and M6 MBONs is at least partially a result of depressed feedforward inhibition from the MVP2 MBON (11; 14). To the best of our knowledge, simultaneous changes in approach and avoidance MBON activity has not yet been observed. A consequence of this coordinated plasticity is that, if plasticity onto one MBON type is blocked, plasticity at the other MBON type should compensate.

#### Prediction 5 – MV model

DANs of both valence modulate plasticity at MBONs of a single valence. In contrast, anatomical and functional experimental data suggest that, in each MB compartment, the DANs and MBONs have opposite valences (16; 39). However, the GAL4 lines used to label DANs in the PAM cluster often include as many as 20-30 cells each, and it has not yet been determined whether all labelled DANs exhibit the same valence preference. Similarly, the valence encoded by MBONs is not always obvious. In (9), for example, it is not clear whether optogenetically activated MBONs biased flies to approach the light stimulus, or to exhibit no-go behaviour that kept them within the light. In the absence of anatomical evidence for the MV model, it may be elaborated so that either each compartment contains DANs of both valences, or *ii*) each compartment contains MBONs of both valences (Supplementary Fig. 6). In larval *Drosophila*, there are several examples of cross-compartmental DANs and MBONs (40), but a full account of the valence encoded by these neurons is yet to be provided.

### Previous mushroom body models

We are not the first to present a MB model that learns different rewards for multiple cues for decision making (19–21). However, these models utilise bounded synapses to prevent reward predictions from growing indefinitely with continued reward experience. Thus, given enough training, these models would not differentiate between two cues that were associated with rewards of the same sign, but different magnitudes. Our model builds upon these studies by incorporating feedback from MBONs to DANs, which allows KC→MBON synapses to accurately encode the reward magnitude and sign with stable fixed points that are reached when the RPE signalled by DANs decays to zero.

### Model limitations

Central to this work is the assumption that the MB has only a single objective: to minimise the RPE. In reality, an organism must satisfy multiple objectives that may be mutually opposed. In *Drosophila*, anatomically segregated DANs in the *γ*-lobe encode water rewards, sugar rewards, and motor activity (7; 8; 17; 22), suggesting that *Drosophilae* do indeed learn to satisfy multiple objectives. Multi-objective optimisation is a challenging problem, and goes beyond the scope of this work. Nevertheless, for many objectives, the principle that accurate predictions aid decision making, which forms the basis of this work, still applies.

For simplicity, our simulations compress all events within a trial to a single point in time, and are therefore unable to address some time-dependent features of learning. For example, activating DANs either before or after cue exposure can induce memories with opposite valences (23; 41). Nor have we addressed the credit assignment problem: how to associate a cue with reinforcement when they do not occur simultaneously. A candidate solution is temporal difference (TD) learning (37; 38), whereby reinforcement information is back-propagated in time to all cues that predict it. While DAN responses in the MB hint at TD learning(36), it is not yet clear how the MB circuity could implement it. An alternative solution is an eligibility trace (38; 42), which enables synaptic weights to be updated upon reinforcement even after presynaptic activity has ceased.

Lastly, our work here addresses memory acquisition, but not memory consolidation, which is supported by distinct circuits within the MB (43). Incorporating memory stabilising mechanisms may help to better align our simulations of genetic interventions with fly behaviour in conditioning experiments.

### Summary

We have developed a model of the MB that goes beyond previous models by incorporating feedback from MBONs to DANs, and shown how such a MB circuit can learn accurate reward predictions through DAN mediated RPE signals. The model provides a basis for understanding a broad range of behavioural experiments, and reveals limitations to learning given the anatomical data currently available from the MB. Those limitations may be overcome with additional connectivity between DANs, MBONs and KCs, which provide five strong predictions from our work.

## Methods

### Experimental paradigm

In all but the last results section, we apply our model to a multi-armed bandit paradigm (38; 44) comprising a sequence of trials, in which the model is forced to choose between a number of cues, each cue being associated with its own reinforcement schedule. In each trial, the reinforcement signal may have either positive valence (reward) or negative valence (punishment, or negative reward), which changes over trials. Initially, the fly is naive to the cue-specific rewards. Thus, in order to reliably choose the most rewarding cue, it must learn, over successive trials, to accurately predict the rewards for each cue. Individual trials comprise three stages in the following order (illustrated in Fig. 1e): *i*) the model is exposed to and computes reward predictions for all cues; *ii*) a choice probability is assigned to each cue using a softmax function (described below), with the largest probability assigned to the cue that predicts the largest reward; *iii*) a single cue is chosen probabilistically, according to the choice probabilities, and the model receives reinforcement with magnitude *r*_+_ (positive reward) or *r*_−_ (negative reward). The fly uses this reinforcement signal to update its prediction of the cue-specific reward.

### Simulations

#### Connectivity and synaptic weights

##### KC→MBON

KCs (K in Fig. 1b) constitute the sensory inputs (described below) in our models. Sensory information is transmitted from the KCs, of which there are *N*_K_, to two MBONs, M_+_ and M_−_, through excitatory, feedforward synapses. For simplicity, we use a subscript x to label the valence of a given sign (e.g. +, for approach MBONs), and use y to label the opposite valence (e.g. in this case, −, for avoidance MBONs). K_*i*_ synapses onto M_±_ with a synaptic weight *w*_±*i*_, which is initialised with *w*_±*i*_ = 0.1*ξ*_±*i*_ for each run of the model, where *ξ*_±*i*_ is a uniform random variable in the range 0-1.

##### KC→DAN

KCs drive excitatory responses in DANs from the PPL1 cluster (29). In our model, we assume that KCs also provide input to appetitive DANs in the PAM cluster. Thus, K_*i*_ drives D_±_ through unmodifiable, excitatory synapses with weights, ***w***_K_ = *γ***1**, where **1** = [1, 1, *…*, 1]^T^ is a vector of ones of length *N*_K_.

##### MBON→DAN

MBONs provide excitatory feedback to their respective DANs (12–14). In both the valence-specific (VS) and mixed-valence (MV) models, M_±_ synapses onto D_∓_ with unit synaptic weight. In the mixed-valence (MV) model, M_±_ also provides inhibitory feedback to D_±_ via an inhibitory interneuron, but we do not model the interneuron explicitly. Thus, we describe the feedback weight simply as *w*_M_ = 1, and specify whether the input is excitatory or inhibitory in the firing rate equation for D_±_ (Eqs. 11 and 12).

#### Inputs and KC sensory representation

Projection neurons from the antennal lobe and optic lobes provide a substantial majority of inputs to Kenyon cells (KCs) in the MB. These inputs carry olfactory and visual information and, together with recurrent inhibition from the anterior paired lateral neuron, drive a sparse representation of sensory information in approximately 5-10% of the KCs (45–47). For simplicity, we bypass the computations performed in nuclei upstream of the KCs, and assign a unique population of 10 KCs to each cue. Thus, for *N*_c_ cues, we simulate *N*_K_ = 10*N*_c_ KCs. Each KC is always activated by its assigned cue, and each active KC, *j*, is given the same firing rate, *k*_*j*_ = 1 Hz. In a subset of simulations used for Fig. 5d, we simulate 2000 KCs, where each KC is assigned to a cue with probability *p* = 0.05, so that 5% of KCs, on average, are active for a given cue. In these simulations, we normalised the total KC firing rates for each cue, *i*, such that 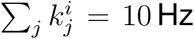. This ensured that the multiplicative effect of KC firing rates on the speed of learning (Eqs. 2 and 5) does not confound the interpretation of our results.

#### MBON firing rates and reward predictions

Neurons are modelled as linear-nonlinear (LN) units that output a firing rate, *y*, equal to the rectified linear sum of their inputs, ***x***:

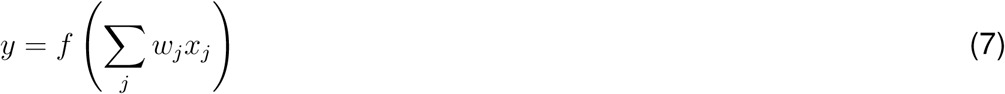

where *f* (*z*) = max(0, *z*) is the rectifying nonlinearity. Eq. 7 can be written more concisely in vector notation: *y* = *f* (***w***^T^***x***), where ***w***^T^ = [*w*_1_, *…*, *w*_*N*_] for *N* presynaptic neurons, and superscript T denotes the transpose. Throughout this text, bold fonts denote vectors.

At the beginning of each trial, MBON firing rates, and thus reward predictions, are computed for each cue. The firing rate, *m*_±_, of MBON M_±_, signals the amount of positive (or negative) reward associated with a given cue, labeled *i*, according to

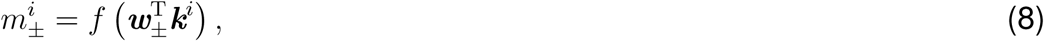

where ***k***^*i*^ is the vector of KC responses to stimulus *i*, and ***w***_±_ are plastic, excitatory synaptic weights. The net reward predicted by sensory cue *i* is then determined by 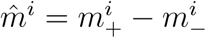.

#### Decision making

In each trial, reward predictions for all cues are compared, and the model is forced to decide which cue should be chosen. Decisions are made probabilistically using a softmax function, *p* (*i*), which specifies the probability of choosing cue *i* as a function of the differences between its reward prediction and the reward predictions of every other cue:

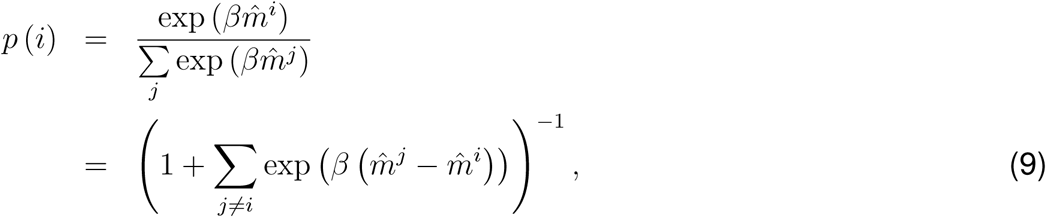

where *β* is a constant (analogous to the inverse temperature in thermodynamics) and modulates the extent to which *p* (*i*) increases or decreases with respect to 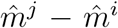. When *β* = 0, choices are independent of the learned valence, and each of the *M* available options are chosen with equal probability, *p* (*i*) = *M*^−1^. When *β* = *∞*, decisions are made deterministically, such that the cue with the highest predicted reward is always chosen. For the multiple-alternative forced choice task, the cue that is ultimately chosen on a given trial is determined by drawing a single, random sample, *ξ*, from a uniform distribution in the range 0-1, and selecting a cue, *q*, such that

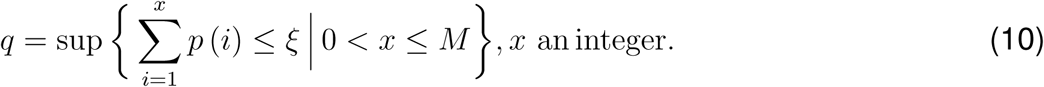

#### DAN firing rates

Once a cue has been chosen, the reward prediction specific to that cue is fed back to the DANs where they are compared against the actual reward, 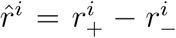, received in that trial, where *r*_±_ is the magnitude of reward signal R_±_. Given the chosen cue, *q*, D_±_ firing rates in the VS models are given by

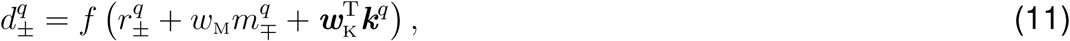

whereas, in the MV model, D_±_ is given by

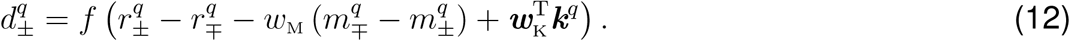

The difference in DAN firing rates yields the RPE for cue *q*:

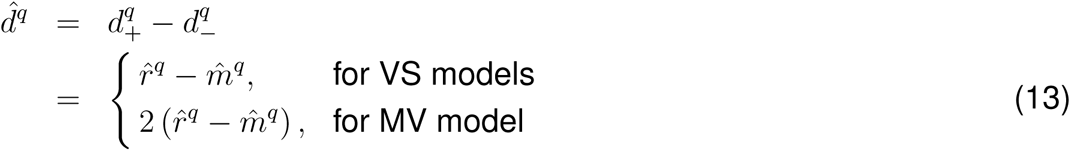

where 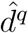 for the MV model is valid when 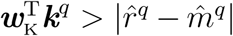. When the inequality is not satisfied, the precise expression for 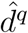 bin the MV model, taking into consideration the non-linear rectification in *d*_+_ and *d*_−_, is

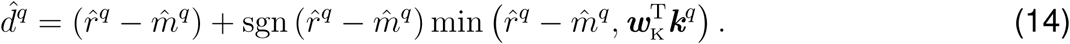

### Synaptic plasticity

We assume that the objective of the MB is to form accurate reward predictions, which minimise RPEs. This objective can be formulated as

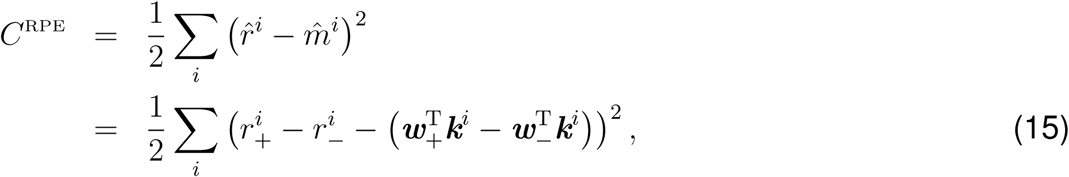

where the sum is over all cues, *i*, 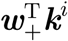 is the firing rate of M_+_, expressed as the weighted input from the KC population response, ***k***^*i*^, through synapses with strength ***w***_+_, and similarly for 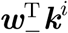. Learning an accurate reward prediction amounts to minimising *C*^RPE^ by modifying the synaptic weights. Assuming that inputs onto approach and avoidance MBONs are modified independently (18), we perform gradient descent on *C*^RPE^ with respect to ***w***_+_ and ***w***_−_ separately. The plasticity rule, 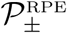, is then defined by the negative gradient:

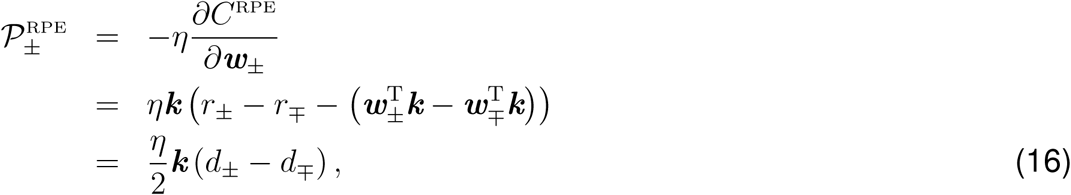

where *η* is the learning rate.

We take the same approach to derive a plasticity rule for the VS model. We start with the cost function

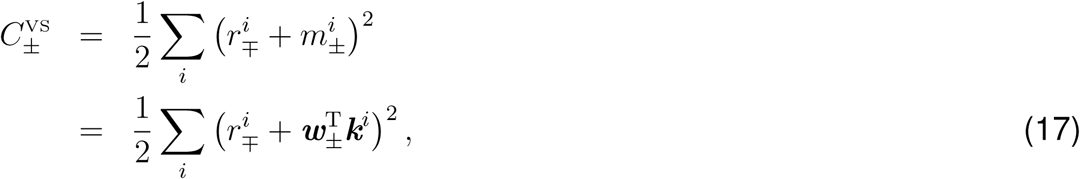

and compute the plasticity rule, 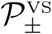, by gradient descent:

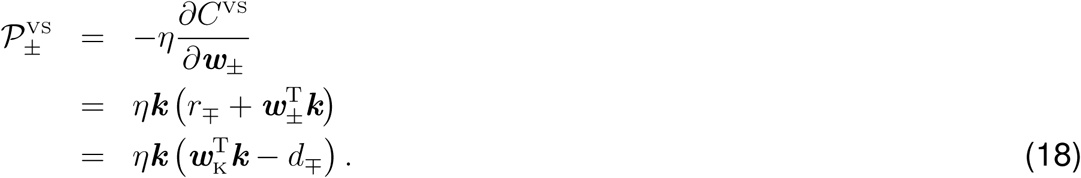

The plasticity rule is in fact only an approximation to gradient descent, and holds true only when the learning rate, *η*, is sufficiently small. Here, sufficiently small means that 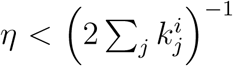 for all cues, *i*, which ensures that learning does not result in unstable oscillations in reward predictions. In our simulations, we set *η* = 2.5 × 10^−2^. The plasticity rule therefore describes the mean drift in synaptic weights over several trials. In the simulations, we use Eqs. 16 and 18 to specify discrete updates to the synaptic weights at the end of each trial, *t*, conditioned on the chosen cue, *q*. Specifically, the update for the VS model is given by

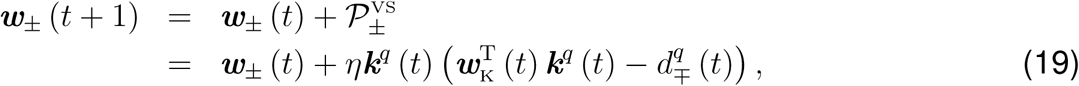

and for the MV model by

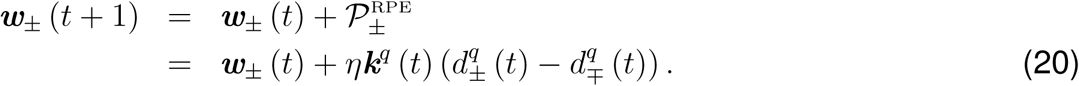

where the superscript *q* specifies the firing rate of each neuron in the presence of cue *q* alone, under the assumption that this cue dominates the neural activity at the point of receiving its corresponding reward signal. The update equation for the VS model with the modified plasticity rule (which we call the VS*λ* model) is

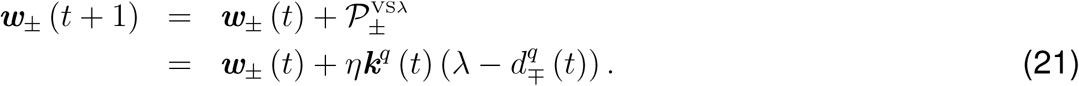

Note that the plasticity rule is not a function of the postsynaptic MBON firing rate (except indirectly through the DAN firing rate). This is possible because a separate plasticity rule exists for synapses impinging on each MBON, negating the need to label the postsynaptic neuron via its firing rate, as would be the case in three-factor Hebbian rules that are typically used in models of reward-modulated learning (26).

### Reward schedule

At the end of each trial, a reinforcement signal specific to sensory cue *i* is provided. Rewards, *r*^*i*^, take continuous values, and are drawn on each trial, *t*, from a normal distribution, *r*^*i*^(*t*) ∼*𝒩* (*µ*_*i*_(*t*), *σ*_R_), with mean *µ*_*i*_(*t*), and standard deviation *σ*_R_. The reward signals that arrive at DANs, R_+_ and R_−_ in Fig. 1D, have amplitudes 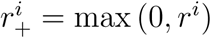 and 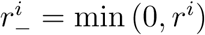, respectively. Over the course of a simulation run, *µ*_*i*_ (*t*) is varied according to a predetermined schedule, and *σ*_R_ is fixed. Thus, at different stages throughout each experiment, the most rewarding cue may switch between the multiple alternatives. Unless otherwise stated, *σ*_R_ = 0.1. The reward schedules were as follows. For Fig. 2 and 3, and Supplementary Fig. 1-4, *µ*_1_ (*t* = 1) = 0, and was held fixed for 20 trials, then underwent a step change of +1 at trials 21, 41, 141, and 161, and a step change of −1 at trials 61, 81, 101, and 121. For Fig. 4 and Supplementary Fig. 5, *µ*_*i*_ (*t*) = *Ag* (*ξ*_*µ*_ (*t*)) + *σ*_R_*ξ*_*σ*_ (*t*), where *ξ*_*µ*_ (*t*) and *ξ*_*σ*_ (*t*) are Gaussian white noise processes with zero mean and unit variance, such that *ξ*_*µ*_ determines the mean reward, and *ξ*_*σ*_ (*t*) determines the additive noise on trial *t*. A low pass filter, *g* (*ξ*_*µ*_) = *F*^−1^{*F*{*ξ*_*µ*_}*F*{*G* (0, *τ*)}}, is applied to *ξ*_*µ*_, where *G* (0, *τ*) is a Gaussian function with unit area, centred on 0, and with standard deviation *τ* = 10 trials, *F*{·} is the Fourier transform, and *F*^−1^{·} is the inverse Fourier transform. Because the Fourier transform method of filtering assumes *ξ*_*µ*_ (1) = *ξ*_*µ*_ (*N*_*t*_ + 1), where *N*_*t*_ is the number of trials, we generate *ξ*_*µ*_ for 250 trials, then delete the first 50 trials after filtering. Finally, the reward amplitude is determined by 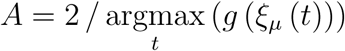.

### Experimental data and model comparisons

The VS*λ* and MV models were compared to experimental data by simulating an often used conditioning protocol. To align with experiments, each simulation utilised the following procedure (Fig. 5a): *i*) in the first stage of training, the model is exposed to a single cue by itself, the CS+, for 10 trials, with rewards drawn from a normal distribution, 𝒩(*µ*, 0.1), where *µ* was chosen according to whether appetitive (*µ* = −1), aversive (*µ* = −1), or neutral (*µ* = 0) conditioning was simulated; *ii*)during the next 10 trials, the model is exposed to a second cue by itself, the CS−, with rewards drawn from a distribution with *µ* = 0 and the same variance as for the CS+; *iii*) the final 2 trials comprise the test stage, in which the model is exposed to both cue 1 and cue 2, as in the MAFC task with two alternatives, with *µ* = 0 for both cues. On each test trial, the model is forced to choose either cue 1 or cue 2, using Eq. 9. We used 10 trials per training stage as, given the parameters for *η* (learning rate) and *β* (inverse temperature), it took this many trials for the mean performance (see below for how performance is measured) across multiple runs of the simulation to plateau at, or near, the maximum possible value. The test was run for only 2 trials as synaptic plasticity was allowed to continue during the test stage, under the assumption that the formation of new CS+ related short term memories (13; 14) might alter the behaviour of flies in the test stage of experiments.

For each simulation, we applied one of many possible additional protocol features, in which neuronal activity was manipulated. We therefore define a protocol as a unique combination of four features:

1. US valence (Fig. 5a): *i*) appetitive (*µ* = 1); *ii*) aversive (*µ* = −1); *iii*) neutral (*µ* = 0).
2. Intervention type (Fig. 5c), which modified the target neuron’s output firing rate from *y*_targ_ to 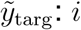) block of neuronal output (e.g. by *shibire*), which was simulated by multiplicatively scaling the manipulated neuron’s firing rate, such that 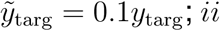) neuronal activation (e.g. by dTrpA1), which was simulated by adding a constant current, such that 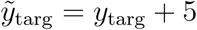.
3. The intervention type was applied following one of four intervention schedules (Fig. 5b): *i*) during the CS+ only; *ii*) throughout training (CS+ and CS−); *iii*) during test only; *iv*) throughout all stages.
4. The target neuron to which the intervention type was applied (Fig. 5c): *i*) M_+_; *ii*) M_−_; *iii*) D_+_; *iv*) or D_−_.

We compared behavioural data from experiments with that of our model for 27 of the 96 possible variations of these four features. These data were obtained from a total of 439 experiments (235 controls with no intervention, 204 experiments with one of the 27 interventions), which were described in 14 published studies (7–13; 22; 23; 27; 30; 31; 33; 34).

Simulations were run in batches of 50, each batch yielding 100 choices from the two test trials. From these choices, we computed a performance index (PI_mod_) given by

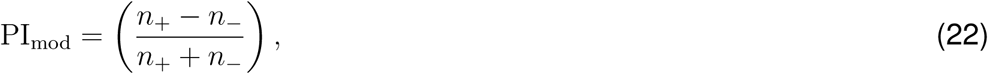

where *n*_+_ is the number of choices for the CS+ and *n*_−_ for the CS−. A distribution of PIs for each protocol was obtained by running 20 such batches. The 439 Pis from the experimental data were extracted by eye from the 14 published papers. These PIs are computed in a similar way as for the model, but where *n*_+_ and *n*_−_ correspond to the number of flies that approached the CS+ and CS−, respectively.

To measure the effect strength of each intervention in both the model and the experiments, we converted PIs into fractions of flies (or model runs) that chose the CS+, *f* = (PI + 1) */*2, then computed a test statistic, Δ_*f*_, which compares *f*_c_ from control to *f*_i_ from intervention experiments, given that the underlying data is binomially distributed, as follows:

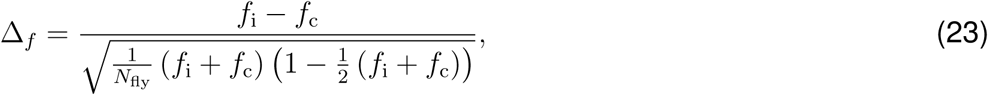

where *N*_fly_ is the number of flies used in that experiment. The binomial distribution adjustment to *f*_i_ − *f*_c_ accounts for the bounded nature of *f* between 0 and 1. As such, for a given absolute difference, *f*_i_ − *f*_c_, Δ_*f*_ is larger when *f*_c_ is near to 1 than when it is near to 0.5. That is, small changes to excellent memory performance imply a stronger effect than small changes to mediocre performance. Because *N*_fly_ was rarely stated in the studies we assessed, we set *N*_fly_ = 50, which is typical for experiments of this nature, and corresponds to the number of runs in each batch of simulations from which a single PI was computed from the model.

To examine the correspondence between PIs from the model and experiments, we fit a weighted linear regression to the experimental versus model Δ_*f*_ data using the MATLAB R2012a function *robustfit*, which computes iteratively reweighted least square fits with a bisquare weighting function. We then computed the Pearson correlation coefficient, *R*, of the weighted data using the weights, ***w***_r_, provided by *robustfit*, according to

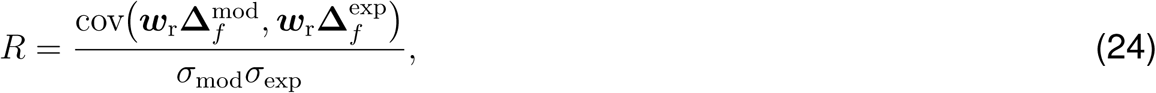

where *σ*_mod_ and *σ*_exp_ are the standard deviations of 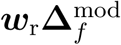 and 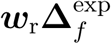, respectively, and bold fonts denote vectors for all data points in either the model or experimental data sets. We determined the probability with which *R* comes from a distribution with zero mean by reshuffling the weighted data

## Data availability

All experimental data was lifted from the data plotted in the cited publications. No additional experimental data was generated in this work. This data is collated and made available at: https:/github.com/BrainsOnBoard/MB_reinforcement_learning/mb_rescorla_wagner.

## Code availability

All of the code that was used for running simulations and analysing data can be found on the Brains on Board GitHub repository: https:/github.com/BrainsOnBoard/MB_reinforcement_learning/mb_rescorla_wagner.

## Acknowledgements

Special thanks to Eleni Vasilaki for helpful discussions and feedback on the mathematical formulations, James Marshal for feedback on the manuscript, and the Waddell and Vogels labs for fruitful discussions on learning in *Drosophila*. Thanks also to the members of the Brains on Board team for their critical feedback at various points throughout this project. This work was funded by the EPSRC (Brains on Board project, grant number EP/P006094/1).s

## Author contributions

J.E.M.B. conceived the model, wrote the code, generated and analysed the data. J.E.M.B. and T.N. conceived the reward schedules and ideal agents to test the models. J.E.M.B., A.P. and T.N. wrote and revised the manuscript.

