## Supplementary Information for "Learning with reward prediction errors in a model of the *Drosophila* mushroom body"

James E. M. Bennett[1], Andrew Philippides[1], Thomas Nowotny[1]

[1]Department of Informatics, University of Sussex, UK

Corresponding author:

James E. M. Bennett

School of Engineering and Informatics

University of Sussex

Chichester 1, Room 002

Falmer

Brighton

BN1 9QJ

### Supplementary Text

In Fig. 5d-e we highlighted an example experimental intervention that revealed substantial differences between the learned behaviour produced by the VS$\lambda$ and the MV models. The intervention comprised aversive conditioning with shibire block of M$+$ (approach MBON) in the training phase during both CS$+$ and CS$-$ exposure. In the following, we describe how the two models produce these different behaviours.

In this example, the VS$\lambda$ model yielded a strong reduction in the number of CS$+$ choices as compared with controls, whereas there was a negligible difference in the choice behaviour of the MV model. There are two factors at play here, which we outline and then backup by analysing the equations for the synaptic weight dynamics. First, inhibitory block of M$_+$ in the MV model amplifies the difference between reward predictions for the CS$+$ and CS$-$, $\hat{m}_{\mathrm{CS+}} - \hat{m}_{\mathrm{CS-}}$, whereas in the VS$\lambda$ model, $\hat{m}_{\mathrm{CS+}} - \hat{m}_{\mathrm{CS-}}$ is the same as in the control. Second, $\hat{m}_{\mathrm{CS+}}$ decays to zero more rapidly in the VS$\lambda$ model than in the MV model during the test phase. Combining this second point with the fact that $\hat{m}_{\mathrm{CS+}} - \hat{m}_{\mathrm{CS-}}$ is smaller in the VS$\lambda$ model implies that $\hat{m}_{\mathrm{CS+}} = \hat{m}_{\mathrm{CS-}}$ occurs much sooner in the VS$\lambda$ model, resulting in a reduced preference for the CS$+$.

To explain the effect of the inhibitory block on the magnitude of $\hat{m}_{\mathrm{CS+}} - \hat{m}_{\mathrm{CS-}}$, we analyse the synaptic weight dynamics, $\dot{\boldsymbol{w}}_{\pm}$, given the reward being provided ($r_+ = 1$, $r_- = 0$) and the inhibition of M$_+$. In the VS$\lambda$ model, the weight dynamics are given by:

$$\begin{aligned}
\dot{\boldsymbol{w}}_{\pm}\Big|^{\mathrm{VS}\lambda} &= \eta\left(\lambda - d_{\mp}\right) \\
\dot{\boldsymbol{w}}_{+}\Big|^{\mathrm{VS}\lambda}_{\mathrm{CS}+} &= \eta\left(\lambda - 0.1m_{+} - \boldsymbol{w}_{\mathrm{K}}\boldsymbol{k}\right) && (1) \\
\dot{\boldsymbol{w}}_{-}\Big|^{\mathrm{VS}\lambda}_{\mathrm{CS}+} &= \eta\left(\lambda - r_{+} - m_{-} - \boldsymbol{w}_{\mathrm{K}}\boldsymbol{k}\right) && (2) \\
\dot{\boldsymbol{w}}_{+}\Big|^{\mathrm{VS}\lambda}_{\mathrm{CS}-} &= \eta\left(\lambda - 0.1m_{+} - \boldsymbol{w}_{\mathrm{K}}\boldsymbol{k}\right) && (3) \\
\dot{\boldsymbol{w}}_{-}\Big|^{\mathrm{VS}\lambda}_{\mathrm{CS}-} &= \eta\left(\lambda - m_{-} - \boldsymbol{w}_{\mathrm{K}}\boldsymbol{k}\right), && (4)
\end{aligned}$$

and for the MV model, we have:

$$\begin{aligned}
\dot{\boldsymbol{w}}_{\pm}\Big|^{\mathrm{MV}} &= \eta\left(d_{+} - d_{-}\right) \\
\dot{\boldsymbol{w}}_{+}\Big|^{\mathrm{MV}}_{\mathrm{CS}+} &= 2\eta\left(r_{+} - \left(0.1m_{+} - m_{-}\right)\right) && (5) \\
\dot{\boldsymbol{w}}_{-}\Big|^{\mathrm{MV}}_{\mathrm{CS}+} &= 2\eta\left(-r_{+} - \left(m_{-} - 0.1m_{+}\right)\right) && (6) \\
\dot{\boldsymbol{w}}_{+}\Big|^{\mathrm{MV}}_{\mathrm{CS}-} &= 2\eta\left(m_{-} - 0.1m_{+}\right) && (7) \\
\dot{\boldsymbol{w}}_{-}\Big|^{\mathrm{MV}}_{\mathrm{CS}-} &= 2\eta\left(0.1m_{+} - m_{-}\right). && (8)
\end{aligned}$$

Given sufficient time for the synaptic weights to stabilise during learning, such that $\dot{\boldsymbol{w}}_{\pm} = 0$, we can write expressions for $m_{+}$ and $m_{-}$, from which we can calculate the relative reward predictions for the CS$+$ and CS$-$, and thereby determine the strength of appetitive or aversive behaviour in the two models. In the VS$\lambda$ model, we obtain for the MBON firing rates:

$$\begin{aligned}
m_{+}^{\mathrm{VS}\lambda}\Big|_{CS+} &= 10c \\
m_{-}^{\mathrm{VS}\lambda}\Big|_{CS+} &= c - r_{+} \\
m_{+}^{\mathrm{VS}\lambda}\Big|_{CS-} &= 10c \\
m_{-}^{\mathrm{VS}\lambda}\Big|_{CS-} &= c,
\end{aligned}$$

where $c = \lambda - \boldsymbol{w}_{\text{K}}\boldsymbol{k}$ is a constant. As such, the difference between CS$+$ and CS$-$ reward predictions is:

$$\begin{aligned} \hat{m}^{\text{VS}\lambda}_{\text{CS}+} - \hat{m}^{\text{VS}\lambda}_{\text{CS}-} &= 10c - (c - r_+) - (10c - c) \\ &= r_+. \end{aligned} \tag{9}$$

That is, the net preference for the CS$+$ over the CS$-$ remains the same, as the reward prediction for both is increased by the same amount due to the inhibition of M$_+$.

In the MV model, however, both DANs process the $r_+$ reinforcement signal. As such, inhibition of M$_+$ influences the firing rates of both M$_+$ and M$_-$ as follows:

$$\begin{aligned} m^{\text{MV}}_{+}\big|_{CS+} &= 10\,(r_+ + m_-) \\ m^{\text{MV}}_{-}\big|_{CS+} &= 0.1m_+ - r_+ \\ m^{\text{MV}}_{+}\big|_{CS-} &= 10m_- \\ m^{\text{MV}}_{-}\big|_{CS-} &= m_-. \end{aligned}$$

and consequently, the difference between CS$_+$ and CS$_-$ reward predictions in the MV model is also affected:

$$\begin{aligned} \hat{m}^{\text{MV}}_{\text{CS}+} - \hat{m}^{\text{MV}}_{\text{CS}-} &= 10\,(r_+ + m_-) - 0.1 \times 10\,(r_+ + m_-) - r_+ - (10m_- - m_-) \\ &= 8r_+. \end{aligned} \tag{10}$$

Thus, approach behaviour towards the CS$+$ is eight times stronger in the MV model than in the VS$\lambda$ model. This difference in CS$+$ preference cannot, by itself, explain the behaviours exhibited by the two models. This is because, in our simulations, behaviour is measured in the test phase across two trials, such that learning after the first trial may affect the behaviour in the second trial. Thus, the rate at which the appetitive memory is forgotten in the test phase must also be considered.

The rate of memory decay during the test phase is given by the difference in the synaptic weight dynamics of $\boldsymbol{w}_+$ and $\boldsymbol{w}_-$, taking into account that M$+$ is no longer receiving inhibitory block, and that $r_+ = 0$. Given that, in both the VS$\lambda$ and MV models, the CS$+$ is preferred over the CS$-$, we assume that the CS$+$ is chosen in the first trial of the test phase in both models (which is almost always true in the simulations). We therefore only need to calculate the decay of the CS$+$ memory between trials 1 and 2. For the VS$\lambda$ model, this is:

$$\begin{aligned}
\dot{\boldsymbol{w}}_{\mathrm{CS+}}^{\mathrm{VS}\lambda} &= \dot{\boldsymbol{w}}_{+}^{\mathrm{VS}\lambda} - \dot{\boldsymbol{w}}_{-}^{\mathrm{VS}\lambda} \\
&= \eta_{\mathrm{VS}\lambda}\boldsymbol{k}\left[(\lambda - m_+ - \boldsymbol{w}_{\mathrm{K}}) - (\lambda - m_- - \boldsymbol{w}_{\mathrm{K}})\right] \\
&= \eta_{\mathrm{VS}\lambda}\boldsymbol{k}\,(m_- - m_+) \\
&= \eta_{\mathrm{VS}\lambda}\boldsymbol{k}\,(c - r_+ - 10c) \\
&= -19\,\eta_{\mathrm{VS}\lambda}\boldsymbol{k},
\end{aligned}$$

where we have substituted the stabilised MBON firing rates, as calculated earlier, and used the following parameters from the simulations: $c = 2$ ($\lambda = 12$, $\boldsymbol{w}_{\mathrm{K}} = 10$) and $r_+ = 1$.

The decay rate of the CS$+$ memory in the MV model is given by

$$\begin{aligned}
\dot{\boldsymbol{w}}_{\mathrm{CS+}}^{\mathrm{MV}} &= \dot{\boldsymbol{w}}_{+}^{\mathrm{MV}} - \dot{\boldsymbol{w}}_{-}^{\mathrm{MV}} \\
&= 2\eta_{\mathrm{MV}}\boldsymbol{k}\left[(m_- - m_+) - (m_- - m_+)\right] \\
&= 4\eta_{\mathrm{MV}}\boldsymbol{k}\,(m_- - m_+) \\
&= 4\eta_{\mathrm{MV}}\boldsymbol{k}\,(m_- - 10\,(r_+ + m_-)) \\
&= -40\,\eta_{\mathrm{MV}}\boldsymbol{k},
\end{aligned}$$

where we have assumed that, before training, $\boldsymbol{w}_{+}^{\mathrm{MV}} \approx \boldsymbol{w}_{-}^{\mathrm{MV}}$, and $\boldsymbol{w}_{-}^{\mathrm{MV}}$ were sufficiently low to have reached zero during training (as was the case in our simulations).

Given that $\eta_{\mathrm{VS}\lambda} = 4\eta_{\mathrm{MV}}$ (see Methods in the main text), we find that the CS$+$ memory in the VS$\lambda$ model decays faster than in the MV model. This result, in combination with the fact that the difference between CS$+$ and CS$-$ reward predictions in the VS$\lambda$ model is smaller than in the MV model, tells us that choice behaviour in the VS$\lambda$ model becomes more random much sooner than in the MV model, resulting in the strong relative decrease in CS$+$ choices as shown in Fig. 5d-e.

#### Supplementary Figures and Tables

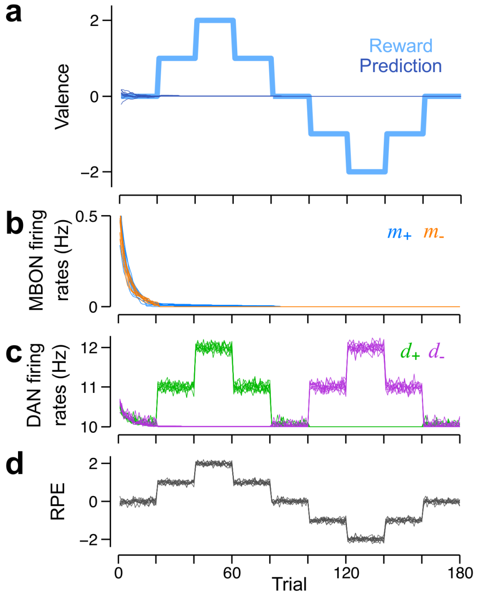


Figure S1: Learning reward predictions in the valence-specific model with the derived plasticity rule given by $\mathcal{P}_{\pm}^{\mathrm{VS}}$ in Eq. 5. **a** Reward schedule (excluding the Gaussian white noise; light blue) and reward predictions (dark blue) from 10 separate runs of the model. Reward predictions in this model necessarily go to zero, because MBON firing rates also necessarily go to zero, as in **b**. **b** Firing rates of the $M_+$ (blue) and the $M_-$ MBONs (orange). **c** Firing rates for the $D_+$ (green) and the $D_-$ (purple) DANs in response to the reward schedule in **a**. **d** Because reward predictions are always zero, RPEs, as given by the difference in firing rates of $D_+$ and $D_-$, are equal to the rewards themselves.

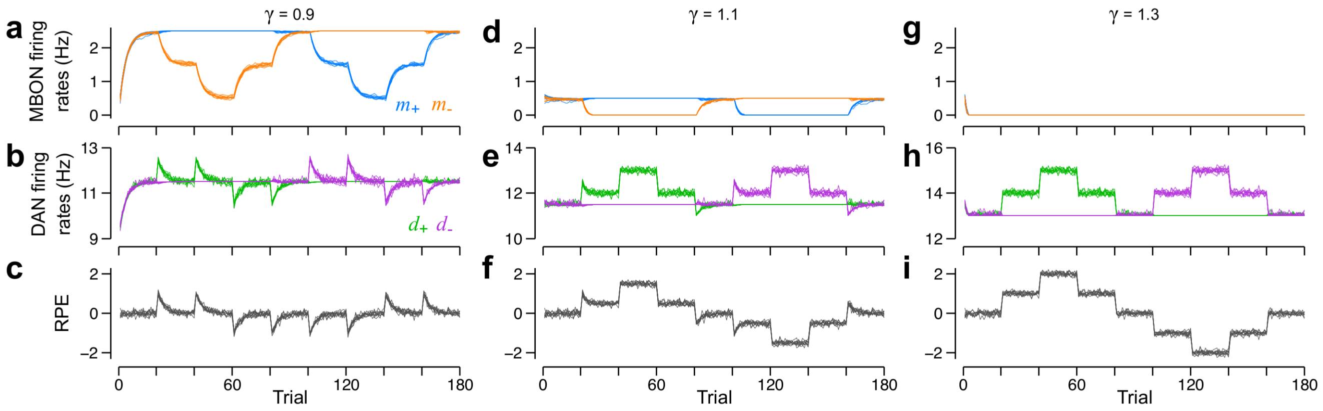


Figure S2: The range of reward predictions that can be learned in the VS$\lambda$ model increases as KC$\rightarrow$DAN synaptic weights are weakened. Each column corresponds to the behaviour of the model with different KC$\rightarrow$DAN synaptic weights, as specified by the $\gamma$ value above each column. The larger the value of $\gamma$, the greater the restriction to the dynamic range of MBON firing rates, and, consequently, the ability of DANs to signal RPEs.

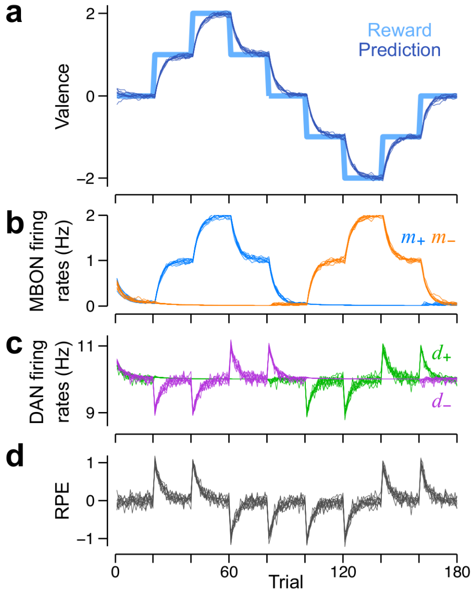


Figure S3: Learning reward predictions in one of the dual valence-specific models (shown in Fig. 3a in the main text, and with the derived plasticity rule, $\mathcal{P}_{\pm}^{\mathrm{VS}}$ in Eq. 5), from which the mixed-valence model is built. **a** Reward schedule (excluding the Gaussian white noise; light blue) and reward predictions (dark blue) from 10 separate runs of the model. **b** Firing rates of the $\mathrm{M}_+$ (blue) and the $\mathrm{M}_-$ MBONs (orange). **c** Firing rates for the $\mathrm{D}_+$ (green) and the $\mathrm{D}_-$ (purple) DANs in response to the reward schedule in **a**. **d** RPEs, as given by the difference in firing rates of $\mathrm{D}_+$ and $\mathrm{D}_-$.

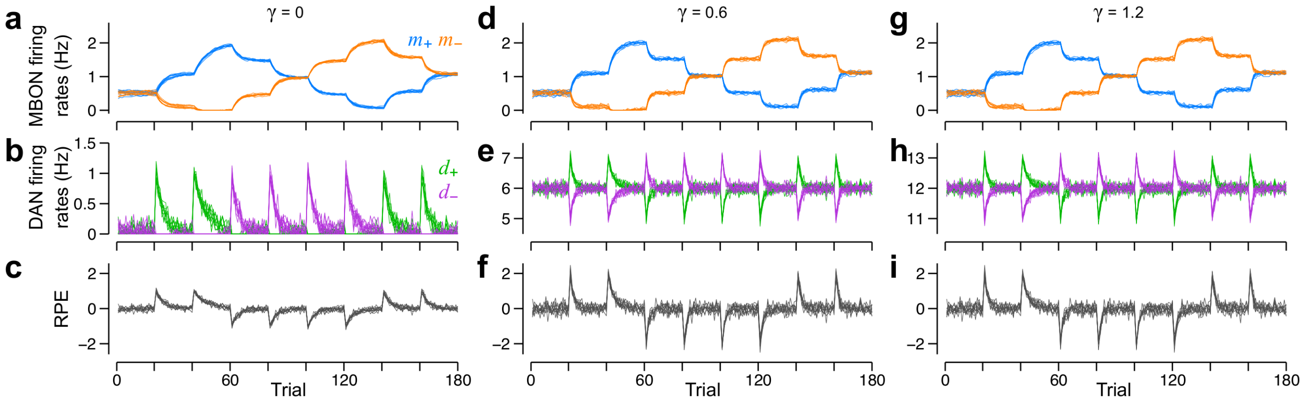


Figure S4: Learning reward predictions in the mixed-valence model. Each column corresponds to the behaviour of the model with different KC→DAN synaptic weights, as specified by the $\gamma$ value above each column. Above a critical value, changing the KC→DAN synaptic weights has no effect on model behaviour. However, when $\gamma$ is chosen such that $\boldsymbol{w}_{\mathrm{K}}^{\mathrm{T}}\boldsymbol{k}$ is less than the RPE magnitude, the dynamic range of DANs is restricted (**a**-**c**). As such, the RPE reported by the difference in firing rates of $D_+$ and $D_-$ is reduced, resulting in slower learning, and RPEs that decay more slowly.

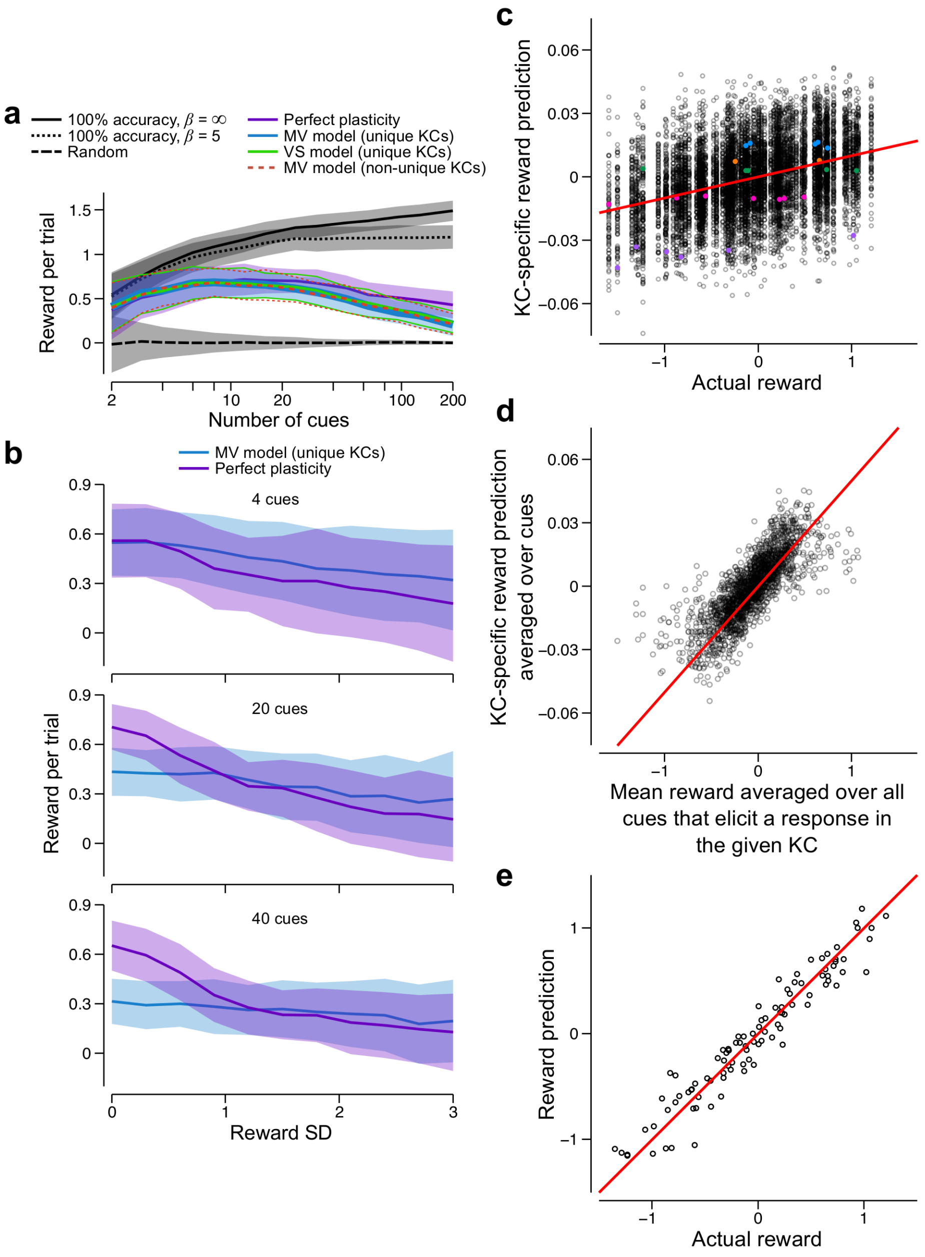


Figure S5: Model performance, as compared with different idealised agents, for a multiple-alternative forced choice task involving 2 or more cues.

**a** The performance of the mixed valence (MV) model, as measured by the mean obtained reward per trial (RPT), and as a function of the number of cues, is shown in blue. As a comparison, we also provide the mean RPT for a perfect agent that always chooses the maximally rewarding cue (black, solid curve) – this emulates an agent that predicts rewards for all cues with 100% accuracy, and is fully deterministic ($\beta = \infty$) – and the mean RPT for an agent that randomly chooses cues with equal probability for each cue (black, long-dashed curve). The mean RPT for the MV model peaks when it has to choose between six cues. This is likely due to the trivial fact that there are more large rewards available when there are more cues, as demonstrated by the monotonic increase in mean RPT for the perfect agent. As a percentage of the RPT for the perfect agent, the performance of the model decreases monotonically with an increasing number of cues.

A fairer comparison against the model (black, dotted curve) is an agent that predicts rewards for all cues with 100% accuracy, yet makes decisions probabilistically using the same softmax function used by the model ($\beta = 5$). This results in only a small reduction in the mean RPT compared to the perfect agent.

*Continued on next page*.

Figure S5: *Continued from previous page.* An even more realistic comparison (purple curve) is an agent that can only update reward predictions for the chosen cue, but otherwise exhibits perfect synaptic plasticity, i.e. the reward prediction is set to be equal to the reward just obtained for a given cue. This substantially diminishes performance, which becomes comparable to the model when the number of cues is fewer than twenty. We conclude that the reduced performance of the model, as compared with the 100% accurate agent, is due to the unavoidable fact that the model's ability to maintain accurate predictions for all cues decreases with the number of cues. This is exacerbated by the slowness of synaptic plasticity, as set by the learning rate.
Finally, we tested a version of the model in which KC responses to different cues may overlap. This model comprised 2000 KCs, similar to estimates for a single hemisphere in *Drosophila* (**?**), of which ~5% (totalling 100 KCs) were randomly selected to respond to each cue (**?**). This means that, when there are 100 cues from which to choose, for example, each KC responds to 5 cues on average. Strikingly, this had negligible effect on the mean RPT obtained by the model (red dashed curve). This may be surprising, as synapses from each KC end up learning the mean reward across those 5 cues (Supplementary Fig. 6a-b). However, because the reward predicted for any one cue is encoded in the synapses from 100 KCs, the weighted input from all KCs to $M_+$ and $M_-$ yields accurate reward predictions. We analyse this feature in greater detail in Supplementary Fig. 5e-g.

**b** A slow learning rate is advantageous when the rewards have a stochastic element. We added zero-mean Gaussian white noise, with SD $\sigma_\xi$, to the same, low-pass filtered reward schedules. The performance of both the MV model and the agent with perfect plasticity decreased with increasing noise. However, the addition of stochastic noise had a much greater impact on the perfect plasticity agent. Any advantage it had over the model was lost for $\sigma_\xi \gtrsim 1.0$. Moreover, the performance of the agent was marginally, though consistently lower than the model in this regime.

**c** Accurate reward predictions (RPs) may be obtained from the ensemble of KC inputs to the MBONs, despite the fact that each KC responds to multiple cues. Here, the mixed valence model is trained on 200 cues. All rewards and RPs are taken form the final trial in the simulation, after learning has converged. Red lines in each panel are lines of unity. Each data point corresponds to the reward for a single cue, and a single KC's contribution – the difference between excitatory currents it elicits in $M_+$ and $M_-$ – to the RP for that cue. The contribution from each KC to a RP is very noisy. This is because each KC can only learn the mean reward over the cues to which it responds. RP contributions for five example KCs are shown in the coloured data points. The RP contribution from each KC is approximately the same for all cues to which it responds. Note that, because each cue elicits a response in 100 KCs on average, the RP contribution from each KC is approximately 1% of the total RP.

**d** Each KC learns the mean reward over all cues to which it responds.

**e** Although the contribution from each KC toward the total RP is inaccurate, the total RP, which is the summed contributions over all responding KCs, is accurate.

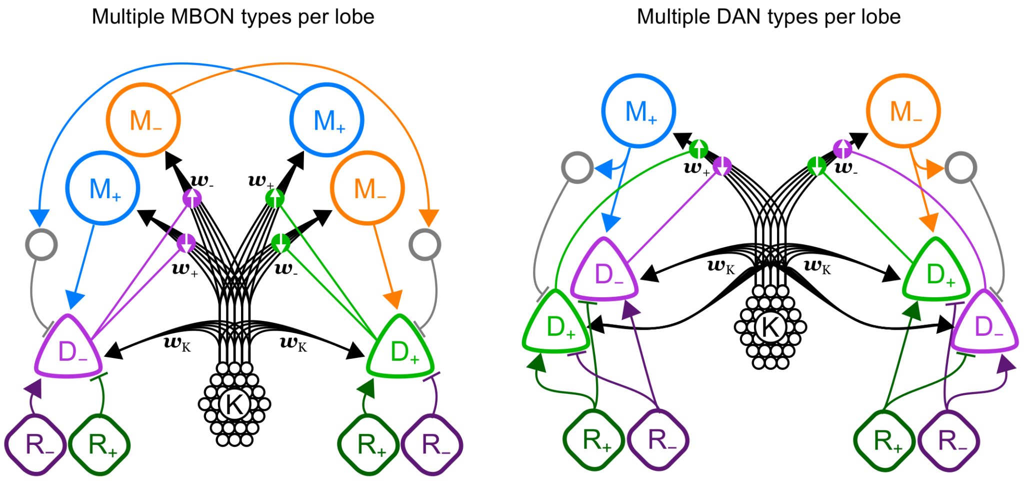


Figure S6: Two elaborated versions of the mixed valence model that avoid the requirement for each DAN to send axons to both mushroom body lobes. *Left*: Each lobe comprises MBONs that bias both approach and avoidance. *Right*: Each lobe comprises DANs that encode both positive and negative reward prediction errors.

| A | B | C | D | E | F | G | H | I | J |
|---|---|---|---|---|---|---|---|---|---|
| **DAN Valence**<br>+1: appetitive<br>-1: aversive | **Effect of DAN on plasticity**<br>+1: potentiation<br>-1: depression | **Reward valence**<br>+1: positive<br>-1: negative | **Input from reward**<br>+1: excitatory<br>-1: inhibitory | **MBON valence**<br>+1: approach<br>-1: avoidance | **MBON outputs**<br>+1: excitatory<br>-1: inhibitory | **IS EFFECTIVE?**<br>B x C x D x E x F > 0 | **IS STABLE?**<br>D + (B x D x F) = 0 | **IS LTP?**<br>B x D > 0 | **CAN LEARN?**<br>G AND H AND I |
| 1 | 1 | 1 | 1 | 1 | 1 | TRUE | FALSE | TRUE | FALSE |
| 1 | 1 | 1 | 1 | 1 | -1 | FALSE | TRUE | TRUE | FALSE |
| 1 | 1 | 1 | 1 | -1 | 1 | FALSE | FALSE | TRUE | FALSE |
| 1 | 1 | 1 | 1 | -1 | -1 | TRUE | TRUE | TRUE | TRUE |
| 1 | 1 | -1 | -1 | 1 | 1 | TRUE | FALSE | FALSE | FALSE |
| 1 | 1 | -1 | -1 | 1 | -1 | FALSE | TRUE | FALSE | FALSE |
| 1 | 1 | -1 | -1 | -1 | 1 | FALSE | FALSE | FALSE | FALSE |
| 1 | 1 | -1 | -1 | -1 | -1 | TRUE | TRUE | FALSE | FALSE |
| 1 | -1 | 1 | 1 | 1 | 1 | FALSE | TRUE | FALSE | FALSE |
| 1 | -1 | 1 | 1 | 1 | -1 | TRUE | FALSE | FALSE | FALSE |
| 1 | -1 | 1 | 1 | -1 | 1 | TRUE | TRUE | FALSE | FALSE |
| 1 | -1 | 1 | 1 | -1 | -1 | FALSE | FALSE | FALSE | FALSE |
| 1 | -1 | -1 | -1 | 1 | 1 | FALSE | TRUE | TRUE | FALSE |
| 1 | -1 | -1 | -1 | 1 | -1 | TRUE | FALSE | TRUE | FALSE |
| 1 | -1 | -1 | -1 | -1 | 1 | TRUE | TRUE | TRUE | TRUE |
| 1 | -1 | -1 | -1 | -1 | -1 | FALSE | FALSE | TRUE | FALSE |
| -1 | 1 | 1 | -1 | 1 | 1 | FALSE | FALSE | FALSE | FALSE |
| -1 | 1 | 1 | -1 | 1 | -1 | TRUE | TRUE | FALSE | FALSE |
| -1 | 1 | 1 | -1 | -1 | 1 | TRUE | FALSE | FALSE | FALSE |
| -1 | 1 | 1 | -1 | -1 | -1 | FALSE | TRUE | FALSE | FALSE |
| -1 | 1 | -1 | 1 | 1 | 1 | FALSE | FALSE | TRUE | FALSE |
| -1 | 1 | -1 | 1 | 1 | -1 | TRUE | TRUE | TRUE | TRUE |
| -1 | 1 | -1 | 1 | -1 | 1 | TRUE | FALSE | TRUE | FALSE |
| -1 | 1 | -1 | 1 | -1 | -1 | FALSE | TRUE | TRUE | FALSE |
| -1 | -1 | 1 | -1 | 1 | 1 | TRUE | TRUE | TRUE | TRUE |
| -1 | -1 | 1 | -1 | 1 | -1 | FALSE | FALSE | TRUE | FALSE |
| -1 | -1 | 1 | -1 | -1 | 1 | FALSE | TRUE | TRUE | FALSE |
| -1 | -1 | 1 | -1 | -1 | -1 | TRUE | FALSE | TRUE | FALSE |
| -1 | -1 | -1 | 1 | 1 | 1 | TRUE | TRUE | FALSE | FALSE |
| -1 | -1 | -1 | 1 | 1 | -1 | FALSE | FALSE | FALSE | FALSE |
| -1 | -1 | -1 | 1 | -1 | 1 | FALSE | TRUE | FALSE | FALSE |
| -1 | -1 | -1 | 1 | -1 | -1 | TRUE | FALSE | FALSE | FALSE |

Table S1: Criteria for unbounded, stable learning of reward predictions. Here, we tabulate the properties of DANs, MBONs, and synaptic plasticity that modulate learning. Columns A-F describe the different properties, each of which takes a value of either $+1$ or $-1$, and each row provides a unique combination those properties. Note that column A is determined by the product of values in columns C and D. Columns G-I determine whether or not each of three criteria, all of which are required for learning, are satisfied by the particular combination of DAN, MBON, and plasticity properties. These criteria are: $i$) that learning induces the expected change in MBON firing rate and thus the expected change in behaviour; $ii$) that learning is stable, so that, for example, the addition of excitatory reward information is offset after learning by the depression of feedback excitation or the potentiation of feedback inhibition. The condition under which each criteria is satisfied is determined by the expression in the second row, which states how the property values in columns B-F must be combined. Column J determines whether or not learning can occur. Only four combinations of properties (highlighted in blue) enable learning, and each one contributes to the MV model.
